# Efferent projections of *Nps*-expressing neurons in the parabrachial region

**DOI:** 10.1101/2023.08.13.553140

**Authors:** Richie Zhang, Dake Huang, Silvia Gasparini, Joel C. Geerling

**Author notes:** Correspondence to: Joel C. Geerling, MD, PhD, PBDB 1320, 169 Newton Rd. Iowa City, IA 52246. Grant sponsors. Iowa Neuroscience Institute (INI) 2021 Accelerator Award (JCG). NIH NINDS K08 (JCG) NS099425. NIH NHLBI T35 (RZ) HL166206.

## Abstract

In the brain, connectivity determines function. Neurons in the parabrachial nucleus (PB) relay diverse information to widespread brain regions, but the connections and functions of PB neurons that express *Nps* (neuropeptide S) remain mysterious. Here, we use Cre-dependent anterograde tracing and whole-brain analysis to map their output connections. While many other PB neurons project ascending axons through the central tegmental tract, NPS axons reach the forebrain via distinct periventricular and ventral pathways. Along the periventricular pathway, NPS axons target the tectal longitudinal column and periaqueductal gray then continue rostrally to target the paraventricular nucleus of the thalamus. Along the ventral pathway, NPS axons blanket much of the hypothalamus but avoid the ventromedial and mammillary nuclei. They also project prominently to the ventral bed nucleus of the stria terminalis, A13 cell group, and magnocellular subparafasciular nucleus. In the hindbrain, NPS axons have fewer descending projections, targeting primarily the superior salivatory nucleus, nucleus of the lateral lemniscus, and periolivary region. Combined with what is known about NPS and its receptor, the output pattern of *Nps*-expressing neurons in the PB region predicts a role in threat response and circadian behavior.

## Introduction

The parabrachial nucleus (PB) is a region of the brainstem that links sensory information to a wide variety of brain regions (Cechetto et al., 1985; Herbert et al., 1990; Saper & Loewy, 1980). Glutamatergic neurons in this region belong to two mutually exclusive macropopulations defined by the transcription factors *Atoh1* and *Lmx1b* (Karthik et al., 2022). These two macropopulations have distinct projection pathways: *Lmx1b* neurons project axons to the amygdala and cerebral cortex, primarily via the central tegmental tract, while *Atoh1* neurons project axons to the hypothalamus and other targets, primarily via a ventral pathway (Karthik et al., 2022). Subpopulations of neurons within these two macropopulations have distinct connections (Huang et al., 2020; Huang et al., 2021; Pauli et al., 2022) and functions, which include thermoregulation, pain, itch, sodium appetite, and conditioned taste aversion (Carter et al., 2015; Gasparini et al., 2019; Geerling et al., 2016; Mu et al., 2017; Nakamura & Morrison, 2008, 2010; Shin et al., 2011).

The *Atoh1* macropopulation includes a subpopulation of neurons that express *Nps* and produce neuropeptide S (NPS; Huang et al., 2022). The PB region in mice contains two clusters of NPS neurons – a rostral group near the lateral lemniscus and a caudal group near the locus coeruleus (Clark et al., 2011; Huang et al., 2022; Liu et al., 2011) – and this pattern is similar in the human brainstem (Adori et al., 2015). In *Nps* Cre-reporter mice, we identified additional NPS neurons extending rostrally alongside the lateral lemniscus, as well as novel populations in the nucleus incertus, lateral habenula, and anterior hypothalamus (Huang et al., 2022).

Little is known about the neurons that produce NPS, but the roles of this peptide and its receptor (NPSR1) have been explored in several genetic and pharmacologic studies. Injecting NPS into the rodent brain increased locomotion, arousal, and body temperature, as well as reducing food intake and producing analgesia (Ensho et al., 2017; Leonard et al., 2008; Li et al., 2009; Peng, Han, et al., 2010; Peng, Zhang, et al., 2010; Rizzi et al., 2008; Smith et al., 2006; Xu et al., 2004). The NPS receptor (NPSR1) is required for its arousal-promoting effect (Duangdao et al., 2009; Fendt et al., 2011; Ruzza et al., 2012; Zhu et al., 2010), and human genetic studies have linked *NPSR1* gain-of-function variants to sleep deprivation, anxiety, asthma, and panic disorders (Domschke et al., 2011; Donner et al., 2010; Gottlieb et al., 2007; Laitinen et al., 2004; Xing et al., 2019).

The connections of NPS neurons in the PB region remain unclear, and mapping their axonal projections would help predict possible functions. Previous studies immunolabeled NPS and identified immunoreactive fibers in the septum, hypothalamus, and midline thalamus (Adori et al., 2016; Clark et al., 2011). However, immunolabeling intensity varied with arousal state (Adori et al., 2016), and it remains unclear which population of NPS neurons projects immunolabeled fibers to each target.

In light of these uncertainties, and given the diverse projection patterns of PB neurons, a comprehensive map of axonal projections from NPS neurons is needed. This information is necessary to elucidate the role of NPS neurons. The goal of this study is to analyze the projection pattern of NPS neurons in the PB region as a framework for future studies exploring their function. To accomplish this goal, we combined Cre-conditional axonal tracing with presynaptic labeling (Carter et al., 2013; Opland et al., 2013) and semi-automated bouton plotting (Grady et al., 2022). Based on their genetic identity as *Atoh1*-derived neurons, we hypothesized that NPS neurons project axons primarily to the hypothalamus, via a ventral pathway and not the central tegmental tract (Huang et al., 2022). Our results support this hypothesis and reveal that NPS neurons also project axons via a robust periventircular pathway and target a novel pattern of output sites.

## Materials and Methods

### Mice

All mice were group-housed in a temperature- and humidity-controlled room on a 12/12-hour light/dark cycle with *ad libitum* access to water and standard rodent chow. Overall, we used n=7 Nps-2A-Cre, n=4 Nps-2A-Cre;R26-LSL-L10GFP, and n=4 C57BL6/J mice ranging in age from 8 to 10 weeks (20–40 g body weight, male and female). Detailed information about each knockin strain is provided in **Table 1**. All mice were hemizygous and maintained on a C57BL6/J background. All experiments were conducted in accordance with the guidelines of the Institutional Animal Care and Use Committee at the University of Iowa.

**Table 1.** Cre driver and reporter mice used in this study.

| Strain | Reference | Source information | Key gene |
| --- | --- | --- | --- |
| *Nps-2A-Cre* | Huang, D., Zhang, R., Gasparini, S., McDonough, M. C., Paradee, W. J., & Geerling, J. C. (2022). Neuropeptide S (NPS) neurons: Parabrachial identity and novel distributions. *J Comp Neurol*, 530(18), 3157-3178. https://doi.org/10.1002/cne.25400 | Available from originating investigators | 2A-Cre inserted immediately after terminal exon in endogenous neuropeptide S gene |
| *R26-LSL-L10GFP* reporter | Krashes, Michael J., et al. "An excitatory paraventricular nucleus to AgRP neuron circuit that drives hunger." *Nature* 507.7491 (2014): 238. | Available from originating investigators<br>http://www.informatics.jax.org/allele/MGI:5559562 | Floxed transcription STOP cassette followed by EGFP/Rpl10 fusion reporter gene under control of the CAG promoter targeted to the Gt(ROSA)26Sor locus |

### Stereotaxic injections

Mice were anesthetized with isoflurane (0.5-2.0%) and placed in a stereotactic frame (Kopf 1900 or 940). We made a midline incision and retracted the skin to expose the skull. Through a pulled glass micropipette (20–30 µm inner diameter), we injected 50–100 nL of AAV8-hEf1a-DIO-synatophysin-mCherry (AAV8-DIO-Syp-mCherry, x 2.5 10^13^ pfu/mL; purchased from Dr. Rachel Neve at the Massachusetts Institute of Technology McGovern Institute for Brain Research Viral Vector Core). In Nps-2A-Cre mice used for formal analysis in the coronal plane, we used separate coordinates to target the rostral or caudal cluster of NPS neurons unilaterally in each case. Rostral coordinates were: 1.90 mm right of midline, 4.80 mm caudal to bregma, and 3.90 mm deep to bregma. Caudal coordinates were: 1.05 mm right of midline, 5.35 mm caudal to bregma, and 4.00 mm deep to bregma. In Nps-2A-Cre;R26-LSL-L10GFP mice used for histology in the sagittal plane, we targeted the rostral PB bilaterally, using the following coordinates: left and right 1.80 mm lateral to midline, 4.80 mm caudal to bregma, and 3.90 mm deep to bregma. Each injection was made over a 5-minute period, using picoliter air puffs through a solenoid valve (Clippard EV 24V DC) pulsed by a Grass stimulator. The pipette was left in place for an additional 3 minutes, then withdrawn slowly before closing the skin with Vetbond (3M). Carprofen (5 mg/kg) was provided prior to incision and again 24 hours later for postoperative analgesia. AAV-injected mice were allowed to survive for 3–5 weeks after surgery to allow for production of Cre-conditional proteins.

### Perfusion and tissue sections

Mice were anesthetized with ketamine (150 mg/kg) and xylazine (15 mg/kg, dissolved with ketamine in sterile 0.9% saline and injected i.p.). They were then perfused transcardially with phosphate-buffered saline (PBS, prepared from 10X stock; P7059, Sigma), followed by 10% formalin-PBS (SF100-20, Fischer Scientific). After perfusion, we removed each brain and fixed them overnight in 10% formalin-PBS. We sectioned each brain into 40 µm-thick coronal slices using a freezing microtome and collected tissue sections into separate, 1-in-3 series. Sections were stored in cryoprotectant solution at -30 °C until further processing.

### Immunohistology

For brightfield labeling (immunohistochemistry), we removed tissue sections from cryoprotectant and rinsed them in PBS. To quench endogenous peroxidase activity, we incubated all tissue sections for 30 minutes in 0.3% hydrogen peroxide (#H325-100, Fisher) in a PBS solution containing 0.25% Triton X-100 (BP151-500, Fisher). After 3 washes in PBS, we loaded sections into a primary antibody solution prepared in PBT-NDS-azide, which is a PBS solution containing 0.25% Triton X-100, 2% normal donkey serum (NDS, 017-000-121, Jackson ImmunoResearch), and 0.05% sodium azide as a preservative (14314, Alfa Aesar). This PBT-NDS-azide solution contained either rabbit anti-dsRed or rat anti-mCherry, and we incubated tissue sections overnight, at room temperature, on a tissue shaker. On the following morning, after 3 PBS washes, we incubated sections for 2 hours in a PBT-NDS-azide solution containing a 1:500 dilution of either donkey anti-rabbit (#711-065-152) or donkey anti-rat (#712-065-153; Jackson ImmunoResearch) secondary antibody. Sections were then washed 3 times and placed for 1 hour in biotin-avidin complex (Vectastain ABC kit PK-6100; Vector), washed 3 times in PBS, and incubated in nickel-diaminobenzidine (NiDAB) solution for 10 minutes. Our stock DAB solution was prepared by adding 100 tablets (#D-4418, Sigma, Saint Louis, MO) into 200 mL ddH2O, then filtering it. We used 1 mL of this DAB stock solution, with 300 µL of 8% nickel chloride (#N54-500, Fisher Chemical) per 6.5 mL PBS. After 10 minutes in NiDAB, we added hydrogen peroxide (0.8 µL of 30% H2O2 per 1 mL PBS-DAB) and swirled sections for 4–5 min until observing dark color change. After two rapid PBS washes, we wet-mounted one or more sections, checked them in a light microscope to ensure optimal staining, and in rare cases replaced sections for up to one additional minute for additional enzymatic staining. Finally, after washing an additional 3 times in PBS, we mounted tissue sections on glass slides (#2575-plus; Brain Research Laboratories). Slides were air-dried, then dehydrated in an ascending series of alcohols and xylenes, coverslipped immediately with Cytoseal 60 (#8310-16 Thermo Scientific), and stored at room temperature until microscope imaging.

For immunofluorescence labeling, we removed tissue sections from cryoprotectant, rinsed them in PBS and loaded them into a PBT-NDS-azide solution containing one or more primary antisera (**Table 2**). We incubated these sections overnight at room temperature on a tissue shaker. The following morning, the sections were washed 3 times in PBS, then incubated for 2 hours at room temperature in a PBT-NDS-azide solution containing species-specific donkey secondary antibodies. These secondary antibodies were conjugated to Cy3, Cy5, Alexa Fluor 488, or biotin (Jackson ImmunoResearch #s 711- 065-152, 711-165-152, 711-175-152, 705-065-147, 705-545-147, 713-545-147, 706-545-148, 706-165-148, 706-065-148, 715-065-15, 712-165-153; each diluted 1:1,000 or 1:500). When biotin was used, sections were washed 3 times in PBS, then incubated for an additional 2 hours in streptavidin-Cy5 (#SA1011; Invitrogen) prepared in PBT-NDS-azide. These sections were then washed 3 times in PBS and mounted on glass slides (#2575-plus; Brain Research Laboratories), then coverslipped using Vectashield with DAPI (Vector Labs). Slides were stored in slide folders at 4 °C until microscope imaging.

**Table 2.** Antisera.

| Antigen | Immunogen Description | Source, host species, RRID | Concentration |
| --- | --- | --- | --- |
| CGRP | Synthetic CGRP peptide. | Abcam, goat polyclonal, cat# ab36001, lot #: GR3330475-13, RRID: AB_725807 | 1:1,000 |
| FoxP2 | *E. coli*-derived recombinant human FoxP2 isoform 1 Ala640-Glu715 | R&D Systems, sheep polyclonal cat# AF5647; lot#: CCUB0109061, RRID: AB_2107133 | 1:3,000 |
| mCherry | Full length mCherry protein | Invitrogen, rat monoclonal, cat# M11217, lot#: VA 293192, lot# W6337875, lot# VH 307508, RRID: AB_2536611 | 1:2,000 |
| GRP | Synthetic (human) bombesin coupled to bovine thyroglobulin with glutaraldehyde. | Immunostar, rabbit polyclonal, cat# 20073, lot# 1420001, RRID: AB_572221 | 1:4,000 |
| AgRP | Human AgRP 83-132 Amide | Phoenix Pharmaceuticals, rabbit polyclonal, cat#: H-003-53, lot#: 01825-2, RRID AB_2313908 | 1:5,000 |
| TH | Native tyrosine hydroxylase from rat pheochromocytoma in sheep host | Millipore, sheep polyclonal, cat#: Ab1542, lot#: 34756625, RRID: AB_90755 | 1:1,000 |
| DsRed | DsRed-express, a variant of *Discosoma* sp. red fluorescent protein | Takara Bio, rabbit polyclonal, cat#: 632496, lot# 1805060, RRID: AB_10013483 | 1:2,000 |
| GFP | Full length green fluorescent protein from the jellyfish *Aequorea victoria* | Invitrogen, chicken polyclonal, cat# A10262, lot# 2591656, lot# 1972783, RRID: AB_2534023 | 1:3,000 |
| TH | Denatured tyrosine hydroxylase from rat pheochromocytoma in rabbit host | Millipore, rabbit polyclonal, AB152, 3199177, RRID: AB_390204 | 1:10,000 |
| NPS | Synthetic NPS peptide | Abcam, rabbit polyclonal, #ab18252, lot: GR198504-1, RRID: AB_776718 | 1:2000 |

### Nissl counterstaining

After whole-slide microscope imaging (described below), all slides were Nissl-counterstained and re-imaged. First, coverslips were removed by soaking slides in xylenes for up to a week. After rehydration, through 1-minute dips in a graded series of alcohols, we rinsed the slides in water and dipped them in a 0.125% thionin solution (Fisher Scientific) for 1 minute. Slides were then rinsed in distilled water until the solution cleared, and then dehydrated in a series of ethanol solutions: 50% EtOH, 70% EtOH, 400mL of 95% EtOH with 10 drops of glacial acetic acid, 95% EtOH, 100% and 100% EtOH. Finally, after the second of two xylene solutions, slides were coverslipped immediately with Cytoseal.

### Microscope imaging and figures

All slides were imaged using an Olympus VS120 slide-scanning microscope. For brightfield images of NiDAB and then Nissl-counterstained sections, we used a 20x UPLSAPO 20x air objective (numerical aperture 0.75) and the extended focal imaging (EFI) to collect and combine in-focus images from 11 focal planes through the tissue. To collect epifluorescence images, we first used a 10x (NA 0.40) objective, then collected additional EFI or multifocal image stacks at 20x in regions of interest. After reviewing whole-slide images in OlyVIA (Olympus), we collected additional EFI or multifocal image stacks at higher magnifications in some regions of interest.

To plot Syp-mCherry-labeled boutons for illustrations from case 5569, we used BoutonNet (Grady et al., 2022). This method uses full-resolution source images (346 nm/pixel OME/TIFF exports from VS-ASW) and combines two separate algorithms to label boutons: an intensity-based “proposal” algorithm, followed by a convolutional neural-network based “confirmation” algorithm. In separate layers in Illustrator, we aligned the PNG output file containing plotted boutons, the NiDAB source image, and a Nissl-counterstained image of the same section. We used the aligned Nissl cytoarchitecture to trace borders, white matter tracts, and ventricles in an additional illustration layer. We inspected each plot to verify accuracy and remove any automated bouton detection atop histological artifacts rather than actual NiDAB-labeled boutons. Finally, we exported the illustration layer and BoutonNet PNG together as a TIF for the final figure layout.

For Figures containing brightfield or fluorescence images, we first used Adobe Photoshop to crop bitmap images from VS-ASW or cellSens (Olympus), adjust brightness and contrast, and combine raw fluorescence data for multicolor combinations. We used Adobe Illustrator to add lettering, trace scale bars atop calibrated lines from OlyVIA (to produce a clean white or black line) and to make all illustrations.

### Whole-brain analysis of Syp-mCherry labeling

To compare the spatial distribution of neurons transduced in the PB region, we first plotted Syp-mCherry-expressing neurons across five rostrocaudal levels from each case with a brain sliced in the coronal plane (approximately -4.7 to -5.6 mm caudal to bregma). We overlaid these injection site plots in Adobe Illustrator and used images of the same sections after Nissl counterstaining to illustrate brainstem borders and major white matter tracts in each section.

Next, we immunolabeled, imaged, and reviewed every tissue section from a 1-in-3 series of sections through the full brain (olfactory bulbs to cervical spinal cord). We began by identifying all regions with Syp-mCherry-labeled boutons in NiDAB-labeled images (without Nissl counterstaining). We then compared each region side-by-side with Nissl-counterstained images of the same sections to identify cytoarchitectural loci. For each region, we scored the density of Syp-mCherry-labeled boutons using a semi-quantitative scale (0–4), and for brain regions with minimal or “trace” labeling, we assigned an intermediate designation (0.5), as in previous work (Huang et al., 2021). Three independent raters (RZ, DH, & JCG) scored every region from every brain and reviewed the results to reach consensus.

#### Nomenclature

We use the term “boutons” to refer to punctate Syp-mCherry labeling. For mouse genes, we used Mouse Genome Informatics (MGI) nomenclature. For mouse proteins, we used common abbreviations from the published literature. For neuroanatomical structures and cell populations, where possible, we used and referred to nomenclature defined in peer-reviewed neuroanatomical literature. In some instances, we used or referred to nomenclature derived from rodent brain atlases (Dong, 2008; Paxinos & Franklin, 2013), with preference to the taxonomy used in the publicly available Allen Brain Mouse Atlas (Wang et al., 2020).

## Results

### Brain-wide distribution of NPS neurons

We began by examining the brain-wide distribution of NPS neurons in parasagittal tissue sections from Nps-2A-Cre;R26-LSL-L10GFP mice (**Figure 1** and **Supplemental Video 1**). The PB region contained the most prominent clusters of NPS neurons, and the sagittal plane of section highlighted the rostrocaudal ribbons of NPS neurons in the lateral habenular nucleus, as well as epitheliod cells running along the central canal of the spinal cord (**Figure 1e**), complementing our previous description in the coronal plane (Huang et al., 2022). Also mathcing our previous description, fewer NPS neurons were scattered in the hindbrain reticular formation, nucleus incertus region, anterior hypothalamus, and medial amygdala.

**FIGURE 1.**
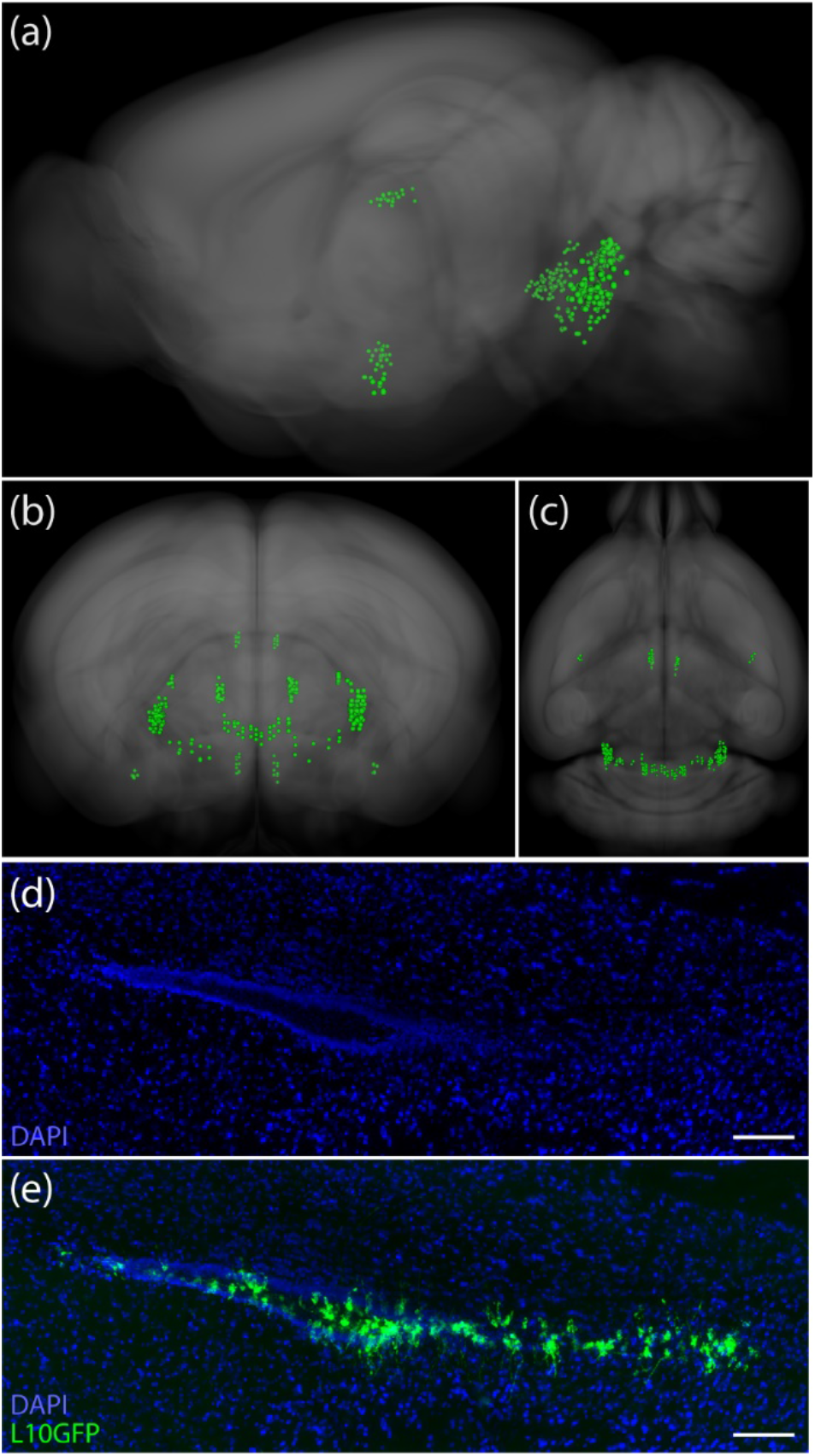
Neurons with L10GFP expression in *Nps* Cre-reporter mice (Nps-2A-Cre;R26-lsl-L10GFP) in a sagittal (a), coronal (b), and horizontal (c) planes. Each dot represents approximately 1–5 neurons. Additional non-neuronal L10GFP-expressing cells were found along the central canal of the spinal cord (d,e). All scalebars are 100 µm.

### NPS-immunoreactive fibers

We immunolabeled NPS and observed fiber labeling in a pattern consistent with a previous report (Clark et al., 2011). The anterior paraventricular thalamic nucleus contained the densest labeling (**Figure 2b**). We also found immunoreactive fibers in the preoptic area (**Figure 2a**), sparse labeling in the dorsal midbrain (**Figure 2c**), moderately dense labeling in the subparaventricular zone (**Figure 2d**), and a dense horizontal band dorsal to the third ventricle (**Figure 2d**). Further caudally in the medial hypothalamus, we found a broad patch of labeling in the DMH and dorsal to it (**Figure 2e**), as well as in the periventricular region alongside the mammillary recess of the third ventricle (**Figure 2f**). All mice had the same overall pattern of fiber labeling, but the intensity of labeling varied between cases.

**FIGURE 2.**
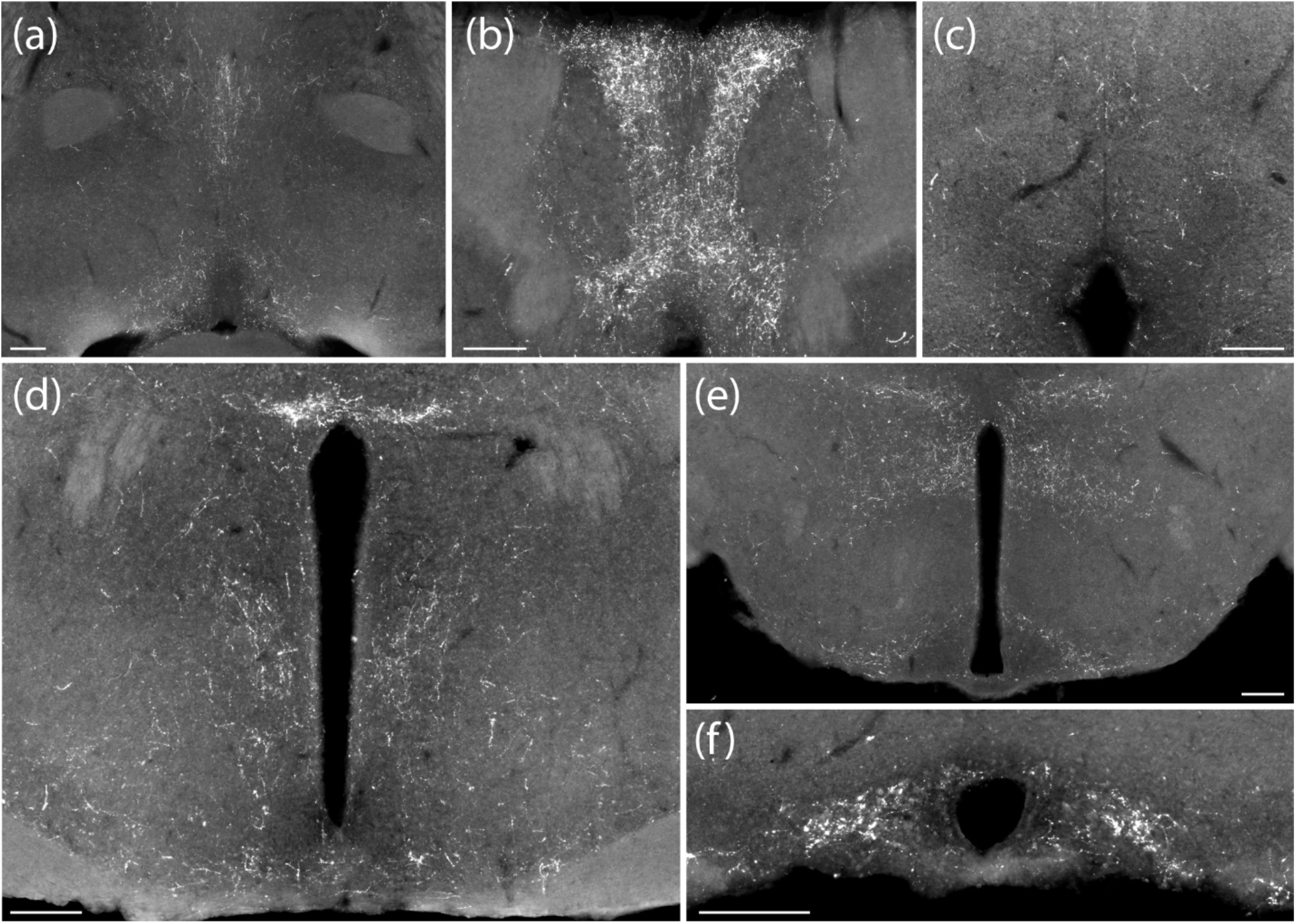
NPS immunofluorescence labeling in the BST (a), PVT (b), PAG (c), PVH (d), caudal hypothalamus (e), and posterior periventricular hypothalamic nucleus (f) varied between cases. Scalebars in every panel are 200 µm.

### Injection sites

Because the origins of NPS-immunoreactive fibers in each region are unclear, we next used Cre-conditional anterograde tracing in the PB region, which contains the densest concentration of *Nps*-expressing neurons. We injected a vector that delivers a Cre-conditional presynaptic marker into the PB of Nps-2A-Cre mice (AAV8-hEf1a-DIO-synaptophysin-mCherry, n=7). We targeted the rostral cluster of PB NPS neurons in three cases (5495, 5568, 5569) and the caudal cluster in four cases. Two caudal injections failed to transduce any neurons, and we analyzed the remaining two cases (5429, 5432).

Rostral-to-caudal plots of each injection site are shown in **Figure 3**. Among the rostral cases, 5568 transduced the most neurons, 5569 transduced an intermediate amount, and 5495 transduced the fewest. Case 5429 mainly transduced neurons in the caudal cluster and transduced fewer neurons in the lateral PB, while case 5432 transduced neurons exclusively in the caudal NPS cluster. We sliced each brain in the coronal plane and used them for brain-wide analysis of Syp-mCherry NiDAB labeling and BoutonNet plots, plus immunofluorescence labeling in select target regions. Rostral-to-caudal images of the full injection site in each case are shown in **Supplementary** Figure 1.

**FIGURE 3.**
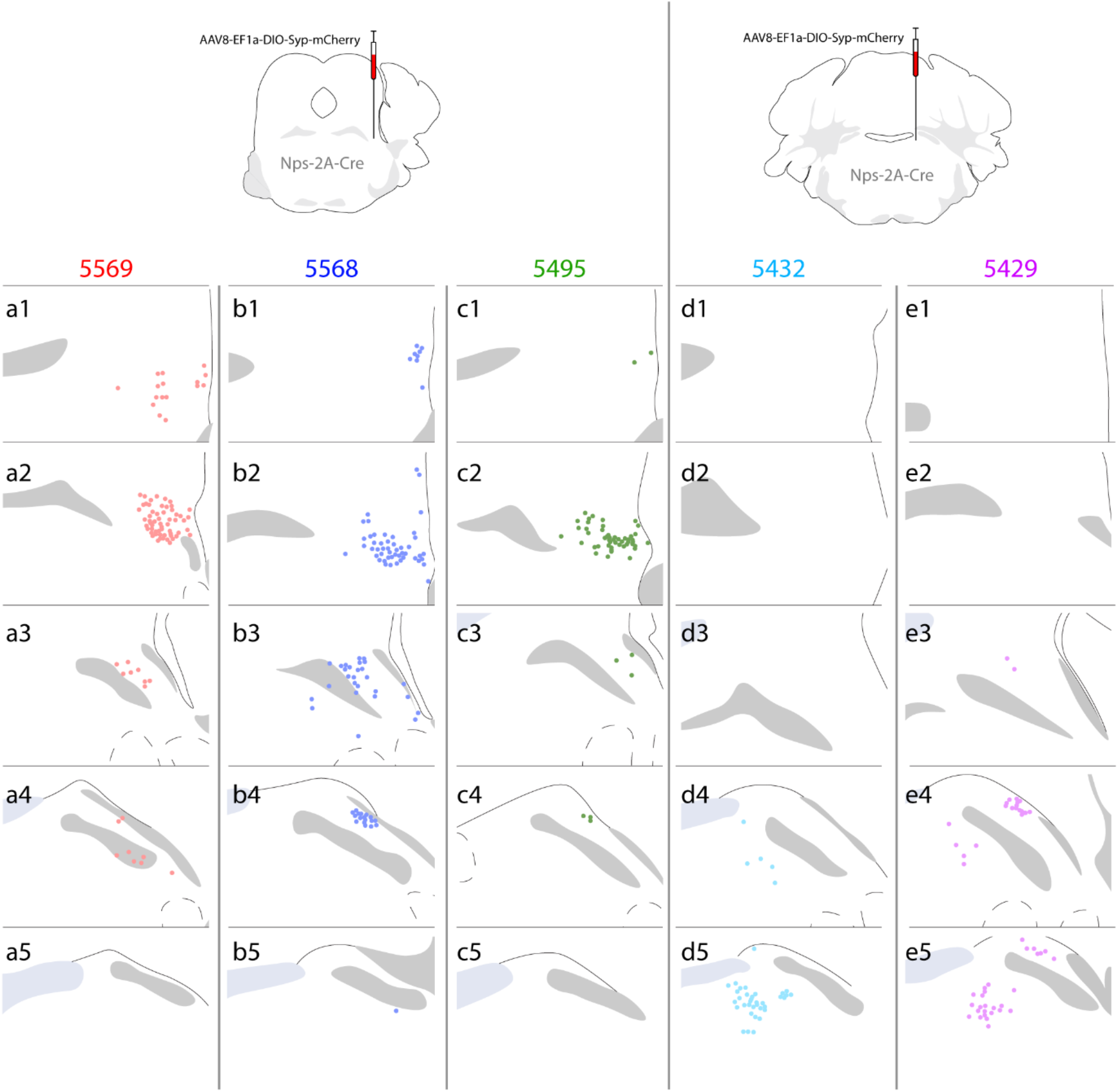
Injection site plots of Syp-mCherry transduced neurons in all Nps-2A-Cre cases. Rostral cases (5569, 5568, 5495) had labeling in the rostral PB with no labeling caudally. In caudal cases (5432, 5429), somatic labeling was primarily located caudally, near the locus coeruleus, and caudal case 5429 had some labeling in the lateral parabrachial nucleus.

As a supplement to our primary analysis in the coronal plane, we injected the same AAV vector into the rostral PB of additional Nps-2A-Cre;R26-LSL-L10GFP mice (n=4). We sliced these four brains in the sagittal plane and used their parasagittal sections for immunofluorescence labeling. An injection site from one of these cases is shown in **Figure 4**.

**FIGURE 4.**
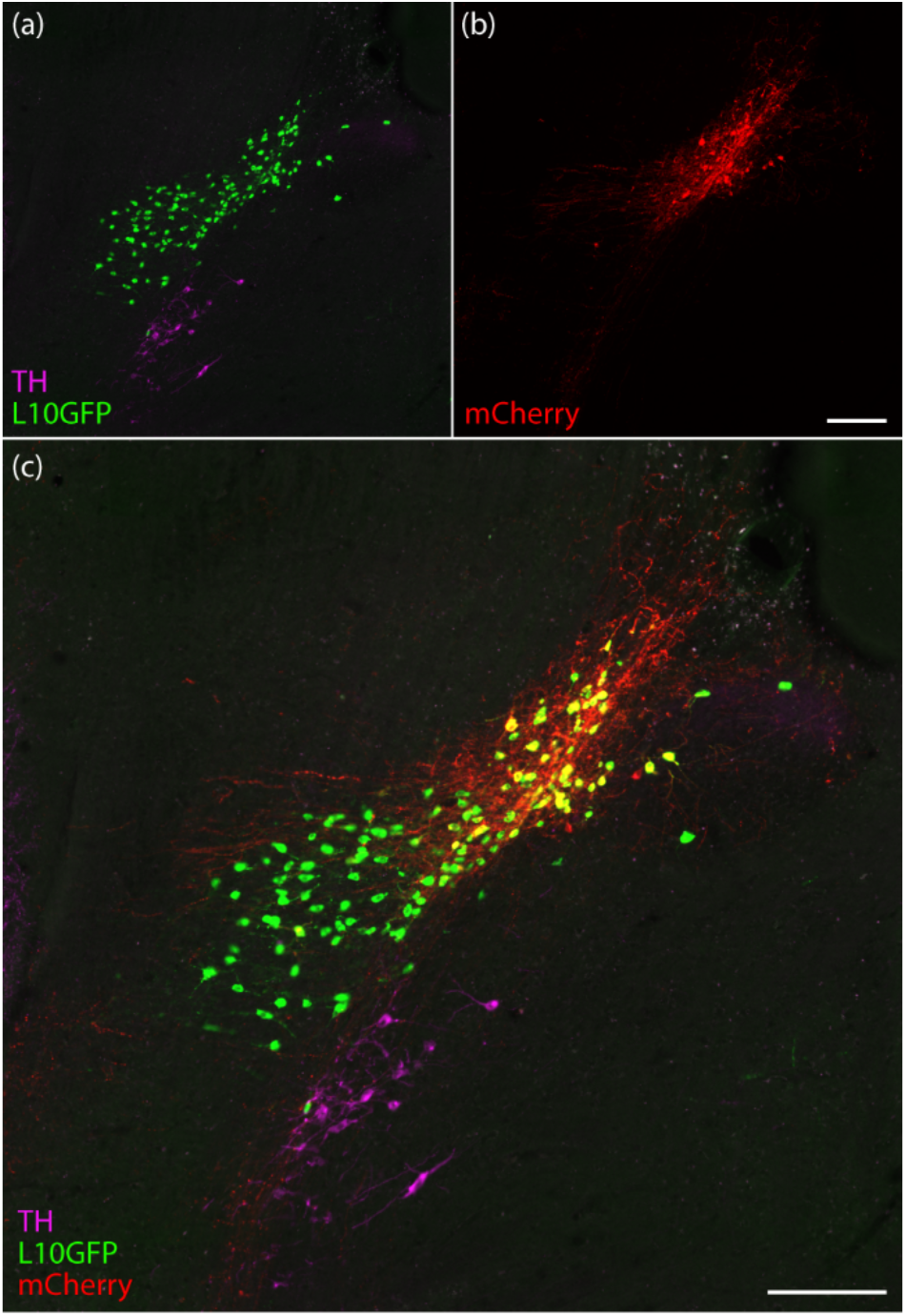
Parasagittal injection site of Cre-conditional Syp-mCherry in case 6730. L10GFP expression reveals cells in the rostral lateral PB with a history of *Nps* expression (green, in a and c). Syp-mCherry expression (red in b) was exclusive to a subset of these neurons (yellow in c). NPS neurons are immediately dorsal and caudal to the noradrenergic A7 group of neurons, which are immunoreactive for tyrosine hydroxlyase (TH; magenta, in a and c). Scalebar in (b) is 200 µm and applies to (a). Scalebar in (c) is 200 µm.

### Brain-wide distribution of Syp-mCherry labeled boutons

We analyzed the brain-wide distribution of Syp-mCherry labeling in each case (**Figure 5**). Overall, cases with injections into the caudal cluster of NPS neurons, near the locus coeruleus, had lighter labeling than those with injections into the larger, rostral cluster of NPS neurons. In rostral cases, nearly every brain region with prominent labeling also contained at least trace labeling in caudal cases. In all cases, labeling ipsilateral to the injection site was stronger. To show the brain-wide distribution of Syp-mCherry-labeled boutons, we performed BoutonNet analysis on tissue sections from a representative case (**Figure 6**).

**FIGURE 5.**
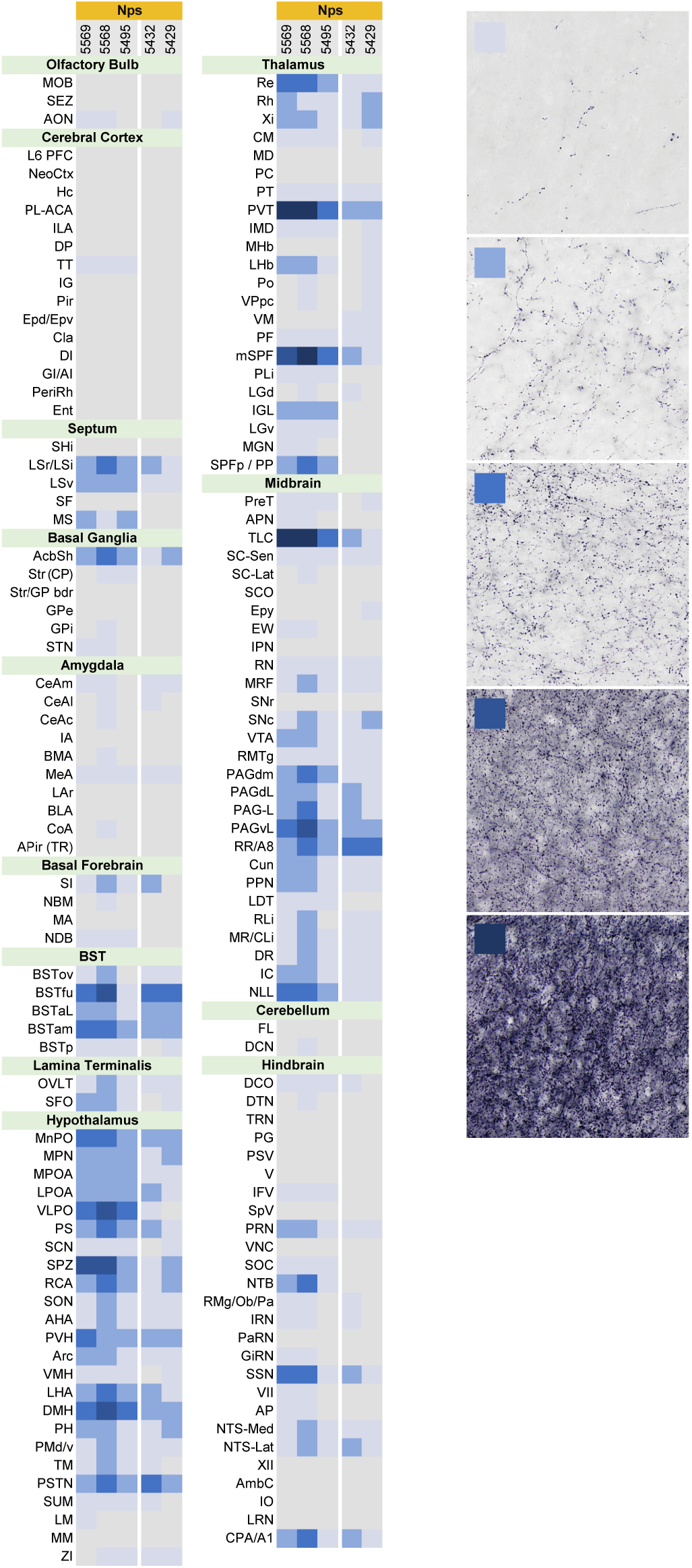
Density of Syp-mCherry labeling in 150 brain regions in each case, grouped by rostral vs. caudal injection site. Light grey indicates absence of labeling and darker shades of blue represent increasing density of labeling. For all abbreviations, see List of Abbreviations.

**FIGURE 6.**
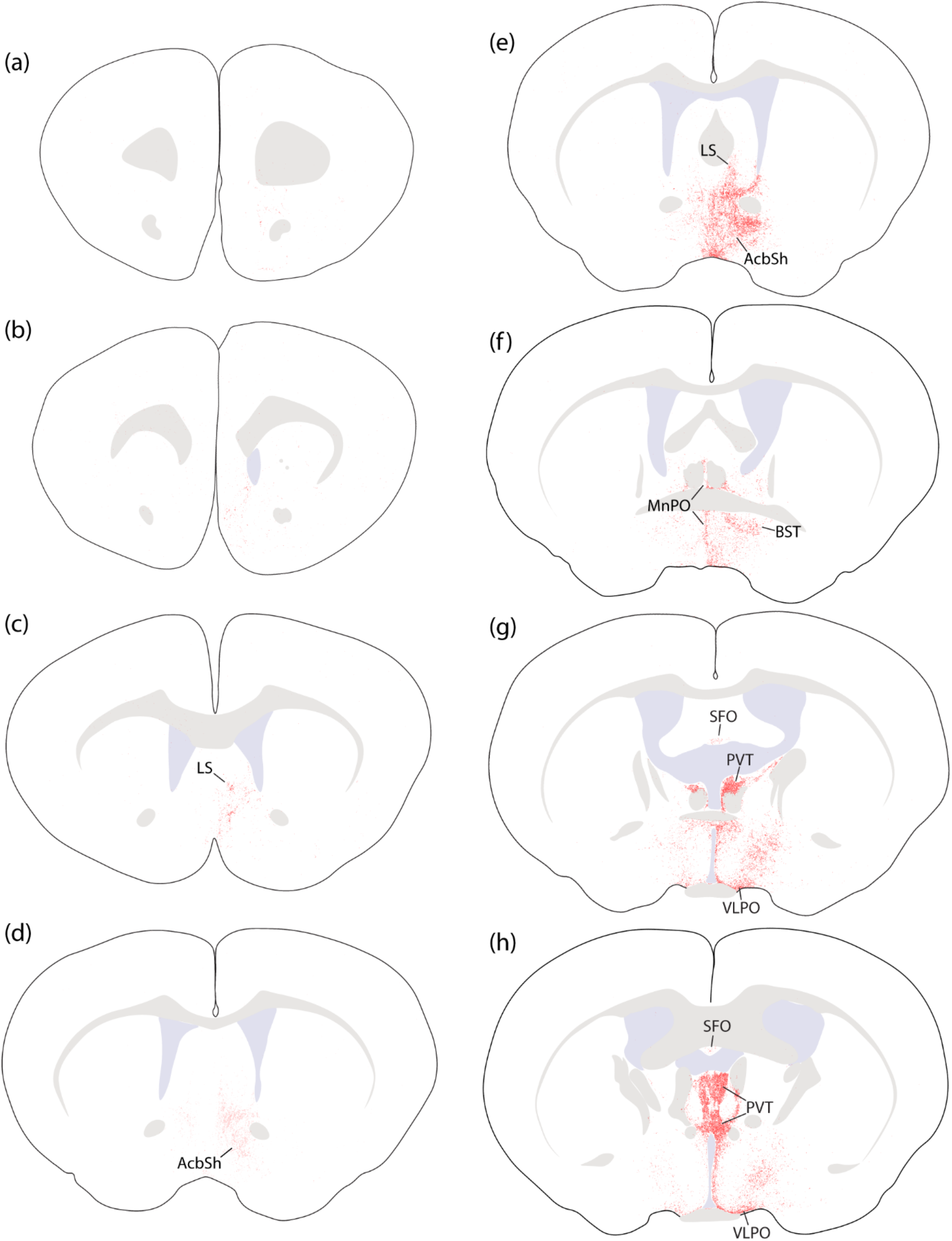
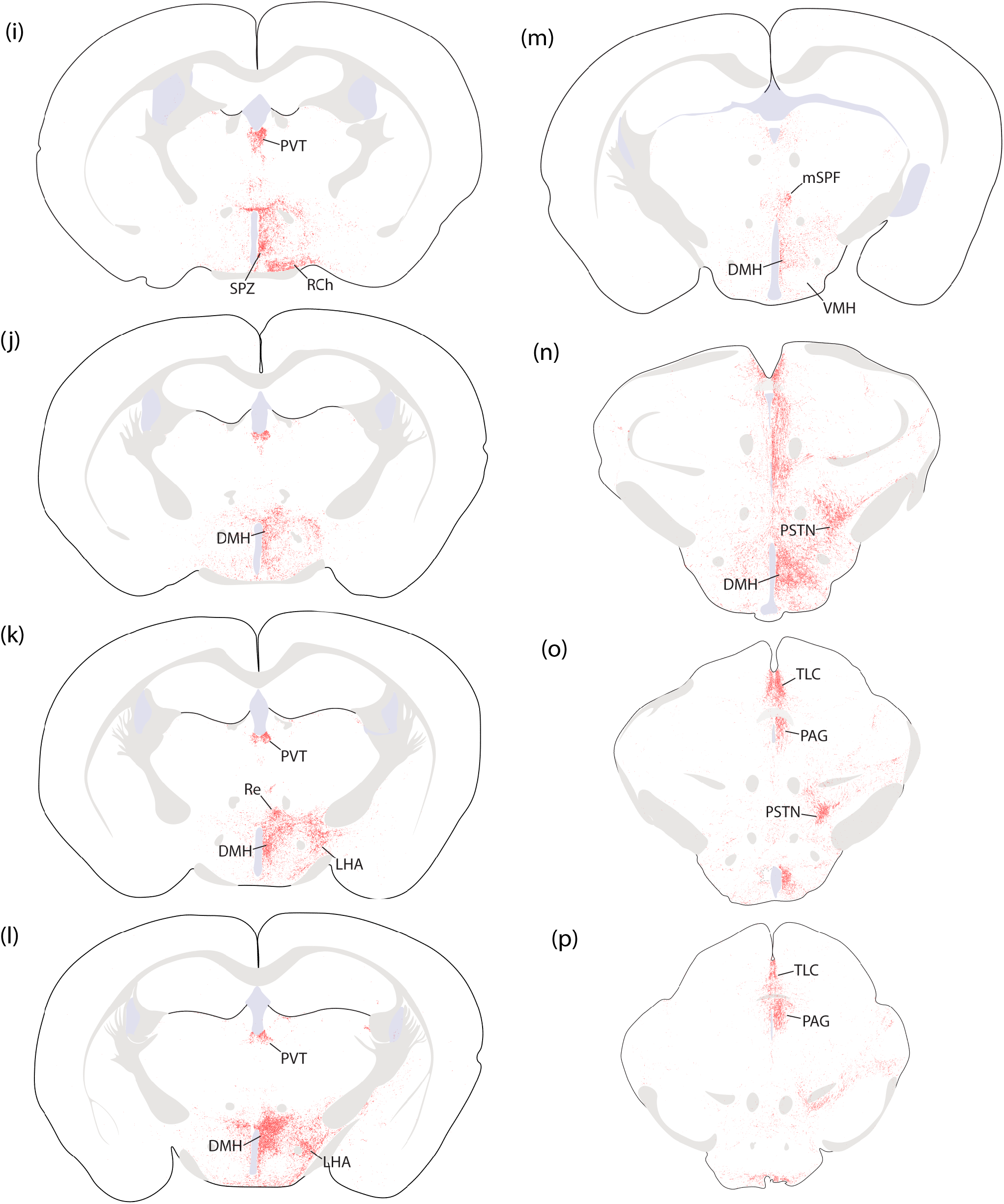
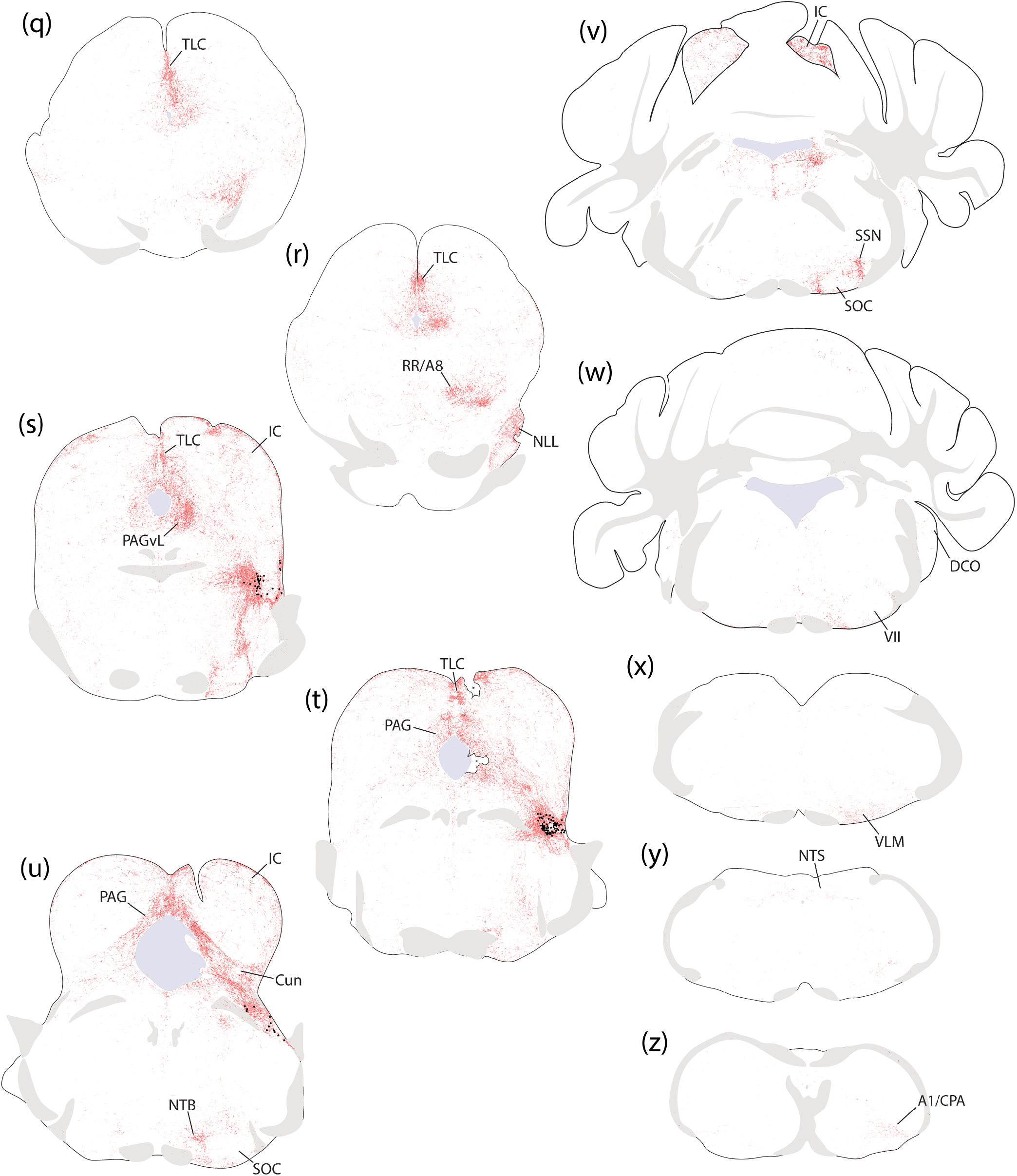
Brain-wide distribution of Syp-mCherry labeling in case 5569. These 26 illustrated sections (a–z) were chosen to best represent the output pattern of NPS neurons in the PB region. Transduced neurons in the injection site are represented by black dots in panels (s–u). All abbreviations are found in List of Abbreviations.

### Cerebral cortex and olfactory bulb

Except for a few, scattered boutons in some cases, we did not find any labeling in the cerebral cortex or olfactory bulb. The anterior olfactory nucleus contained trace labeling in several cases, and the tenia tecta contained trace labeling only in rostral cases.

### Septum

The lateral septum contained labeling in all cases. This labeling concentrated in its rostral and ventral subregions, with virtually no labeling dorsally (**Figure 6c–d**). In rostral cases, the medial septal nucleus also contained light labeling, but no other region of the septum contained labeling in other cases.

### Basal ganglia

In every case, we found light labeling in the caudal nucleus accumbens shell, where it merges with the bed nucleus of the stria terminalis (**Figure 6d-e**). Rostral to this, the nucleus accumbens shell contained no more than trace labeling (**Figure 6c**). Trace labeling in the striatum, globus pallidus, and subthalamic nucleus was an inconsistent finding in rostral cases.

### Amygdala

The amygdala received very little input. The central nucleus of the amygdala contained trace Syp-mCherry labeling in some cases, and the medial amygdalar nucleus had trace labeling in all cases (**Figure 6l**). Only case 5568 had trace labeling in the basomedial nucleus and cortical amygdalar area.

### Basal forebrain

Most basal forebrain nuclei received no input. Some cases had light labeling in the substantia innominata or trace labeling in the diagonal band (**Figure 6c-h**).

### Bed nucleus of the stria terminalis

All cases had dense labeling in the bed nucleus of the stria terminalis (BST). Labeling was densest ventral to the anterior commissure (**Figure 6e-f**), and lighter labeling covered a larger expanse of the anterolateral and anteromedial BST while largely avoiding the oval subnucleus and posterior subnuclei. We immunolabeled AgRP and CGRP to contextualize the anterograde terminal field in this interesting area. Syp-mCherry-labeled boutons overlaid the focal cluster of CGRP-labeled axons that identify the fusiform BST subnucleus, and this medium-density labeling extended medially through ventral BST subnuclei that receive dense AgRP-immunoreactive fibers. In contrast, the oval BST subnucleus, which contains denser CGRP labeling, had significantly less Syp-mCherry labeling (**Figure 7**).

**FIGURE 7.**
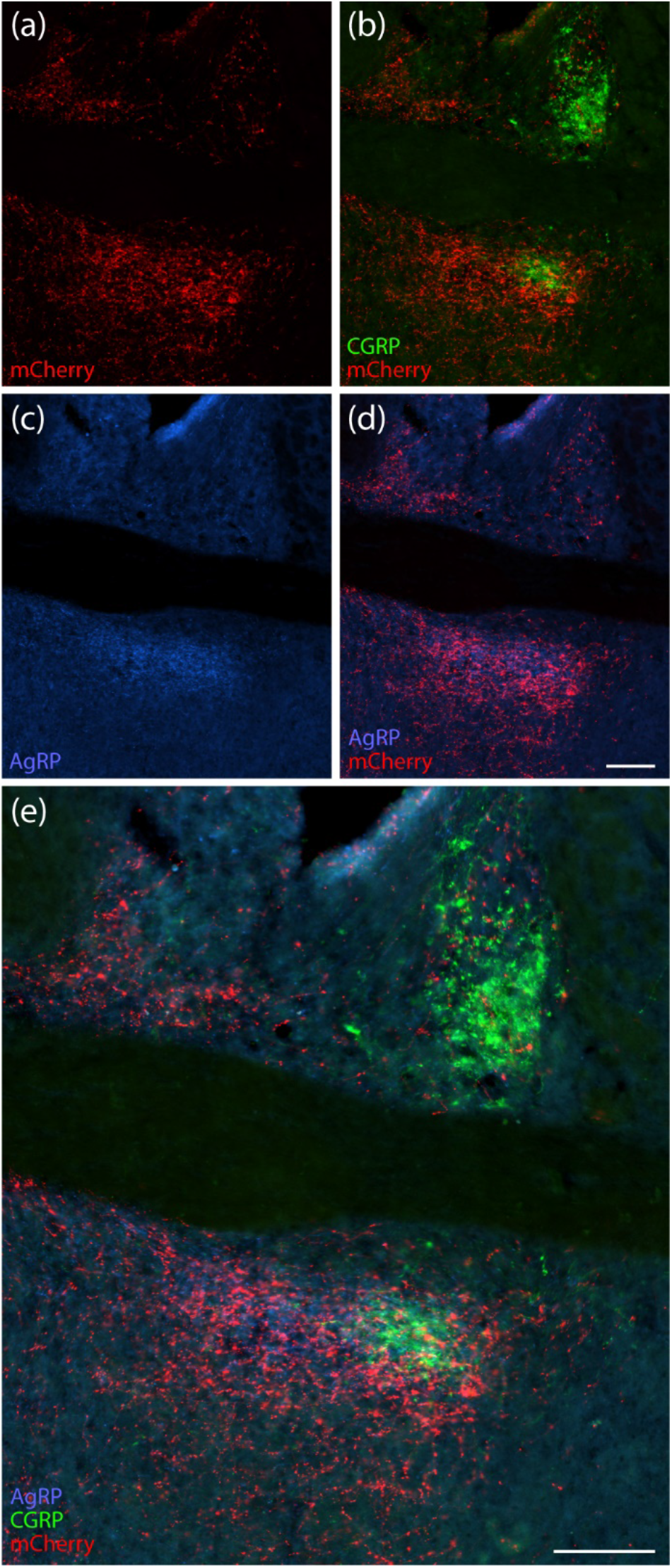
In the ventral BST, Syp-mCherry immunofluorescence (red, a) partly overlapped CGRP-immunoreactive fibers (green, b) and extensively overlapped AgRP-immunoreactive fibers (blue, c–e). AgRP, CGRP, and mCherry combined immunolabeling in panel (e). All sections are from case 5568. All scalebars are 200 µm. Scalebar in (d) also applies to (a–c).

### Lamina terminalis

Both circumventricular organs of the lamina terminalis, the subfornical organ (SFO, Figure 4g) and vascular organ of the lamina terminalis (OVLT), had light labeling in several cases.

In-between them, the median preoptic nucleus contained moderate labeling in each case (**Figure 7**).

### Hypothalamus

In every case, the hypothalamus received more labeling overall than any other brain region (**Figure 5g-o**). However, this labeling was not uniform across the hypothalamus. Syp-mCherry labeling blanketed the medial hypothalamus, except for the mammillary, ventromedial, and suprachiasmatic nuclei, which contained virtually no labeling. Also, a band extending dorsolaterally from the medial preoptic nucleus through the posterior BST received relatively less labeling than surrounding parts of the anterior hypothalamus.

Near the third ventricle and dorsal to the suprachiasmatic nucleus, we found a dense patch of Syp-mCherry labeling in a focal subregion of the subparaventricular zone (SPZ), in-between TH-immunoreactive neurons and axons in the PVH and GRP-immunoreactive fibers projecting dorsally from the SCN (**Figures 6i** & **8**). The paraventricular hypothalamic nucleus (PVH) itself contained only light labeling, but a thin, horizonal band immediately dorsal to the ventral third ventricle contained denser labeling (**Figure 8**).

**FIGURE 8.**
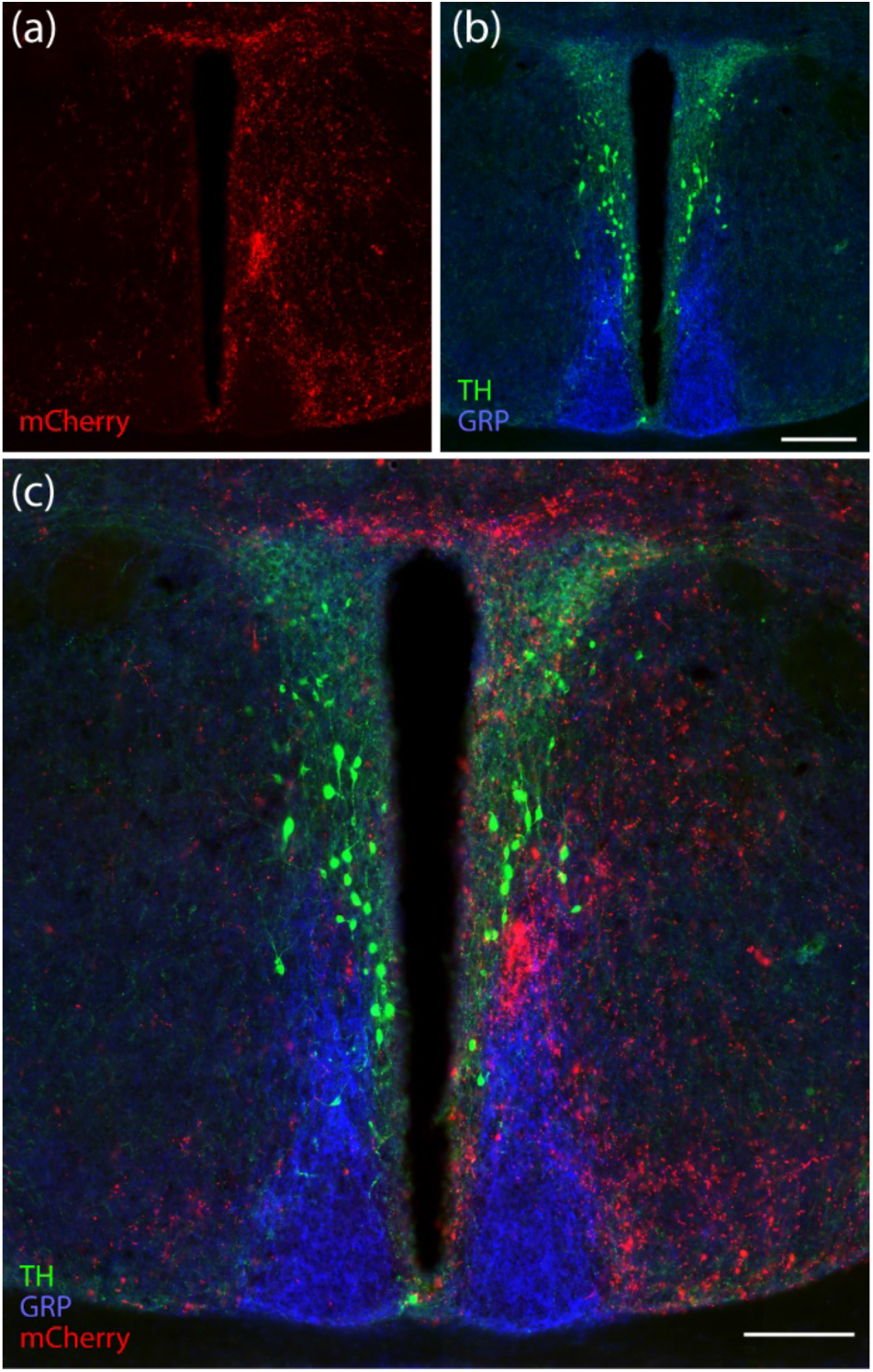
Dense Syp-mCherry immunofluorescence (red, a) lies immediately ventral to TH-immunoreactive neurons and fibers in the paraventricular hypothalamic nucleus (green) and immediately dorsal to GRP-immunoreactive fibers in the SPZ (b–c). Combined mCherry, TH, and GFP labeling is shown in panel (c; case 5569). This level of the anterior hypothalamus also contained dense Syp-mCherry labeling in a band dorsal to the third ventricle. Both scalebars are 200 µm. Scalebar in (b) also applies to (a).

The dorsomedial hypothalamic nucleus (DMH) contained the densest labeling in the hypothalamus, and the parasubthalamic nucleus (PSTN) also contained dense labeling. Labeling in the rest of the lateral hypothalamic area and posterior hypothalamus was less dense. The dense blanket of labeling that covered the DMH extended dorsally to encompass the A13 cluster of catecholaminergic-GABAergic neurons (Negishi et al., 2020) in the zona incerta at the dorsal margin of the hypothalamus (**Figure 9**).

**Figure 9.**
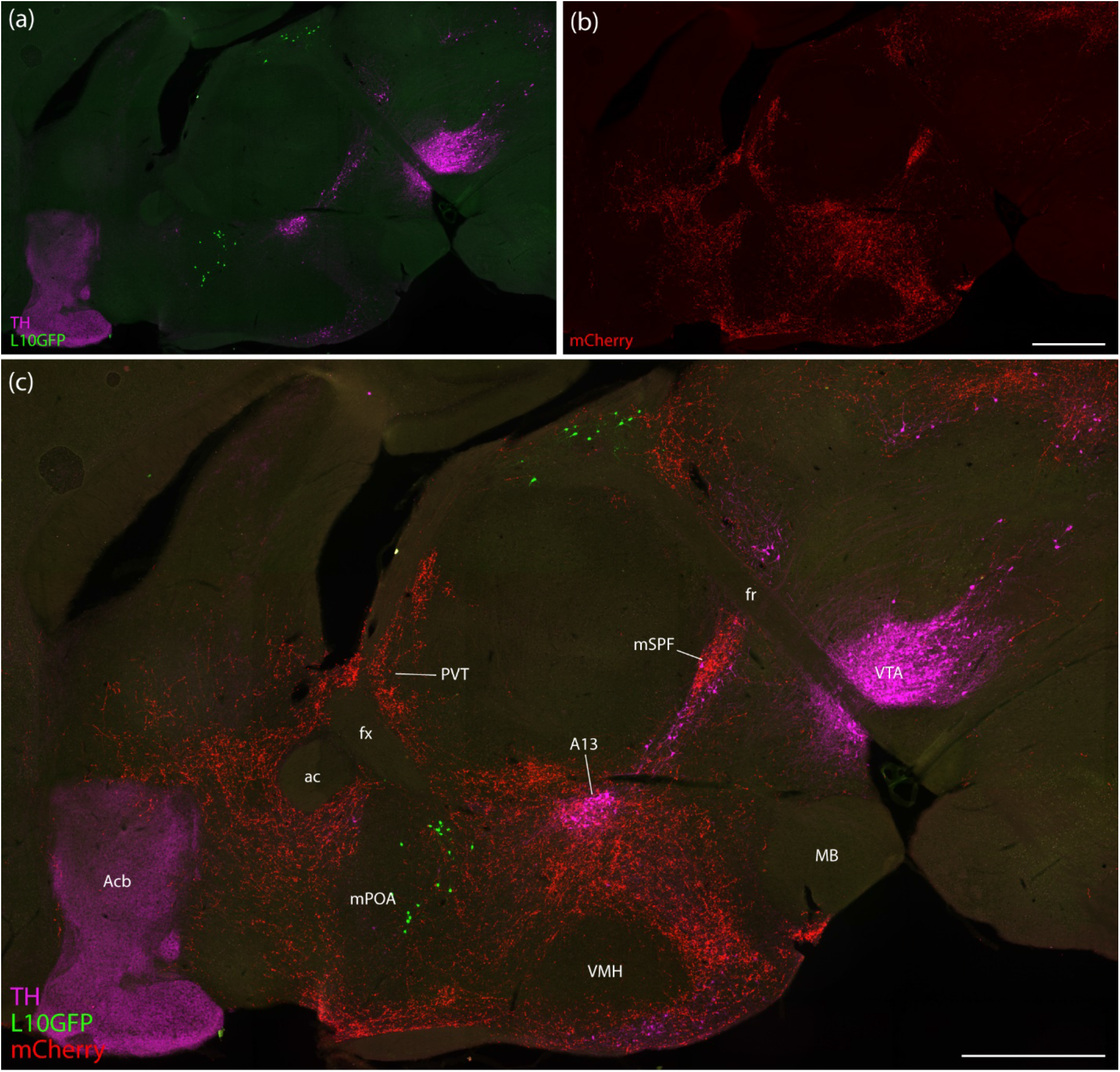
Syp-mCherry labeling (red, b–c) in a parasagittal section though the thalamus and hypothalamus (case 6730). L10GFP Cre-reporter for *Nps* (green; a, c) identifies NPS neurons in the anterior hypothalamic nucleus and lateral habenula, while TH immunolabeling identifies catecholaminergic neurons and fibers (magenta; a, c). Anterogradely labeled axons produced dense concentrations in the paraventricular thalamic nucleus (PVT), magnocellular subparafasicular nucleus (mSPF), A13 group, and several hypothalamic subregions, while most of the thalamus and much of the mPOA, VMH, and MB lacked labeling. For remaining abbreviations, see List of Abbreviations. Scalebars are 800 µm and the scalebar in (b) also applies to (a).

### Thalamus

The median thalamus contained some of the densest labeling in the brain. Every case had dense labeling in the paraventricular nucleus of the thalamus (PVT). Profuse labeling enveloped FoxP2-immunoreactive neurons in the anterior PVT and extended laterally to surround those in the paratenial nucleus (**Figure 10**). This dense terminal field extended caudally through the full PVT. The reunions, rhomboid, xiphoid, and lateral habenular nuclei contained modest labeling.

**Figure 10.**
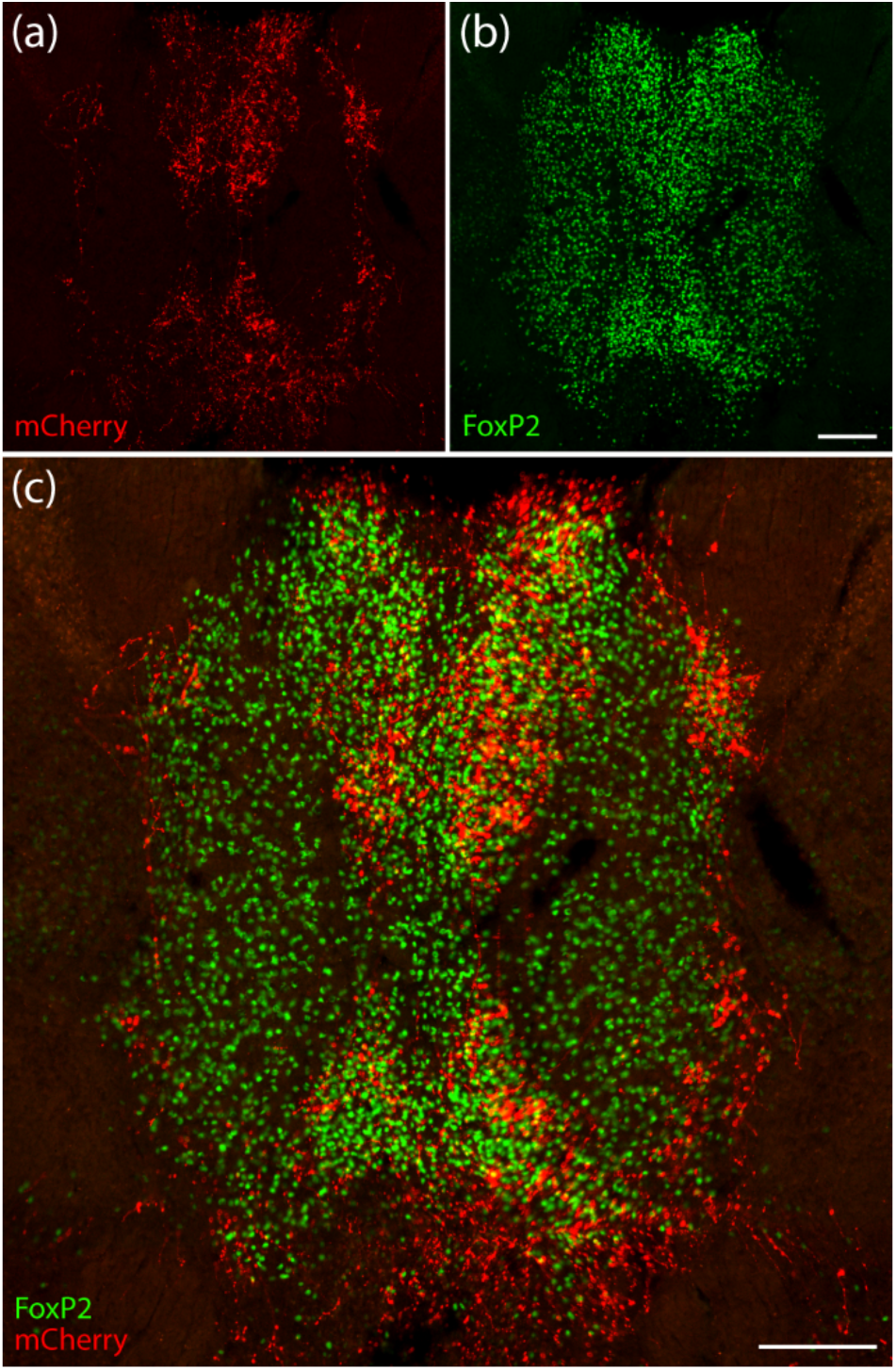
In the thalamus, dense Syp-mCherry labeling (red, a) concentrated in the anterior PVT and arced laterally around the paratenial nucleus (case 5495). These thalamic nuclei contained strong nuclear immunoreactivity for the transcription factor FoxP2 (green, b–c). Scalebar in (b) applies to (a) and is 200 µm. Scalebar in (c) is 200 µm.

We also found a small, dense patch of labeling in the caudal, ventral thalamus. The density of labeling in this small, round spot was similar to the density of Syp-mCherry labeling in the PVT. Syp-mCherry labeling here avoided neighboring catecholamine neurons and instead targeted a small spot referred to as the magnocellular subparafasciular nucleus (mSPF; **Figure 6m**, **Figure 9c**). In contrast, the parafasicular thalamic nucleus had sparse labeling, and the remaining subparafasicular region had light labeling that extended laterally toward the geniculate nuclei and peripeduncular nucleus.

In the caudal, lateral thalamus, we found sparse labeling in the intergeniculate leaflet, ventral lateral geniculate nucleus, and a region referred to as the posterior limitans nucleus (Paxinos & Franklin, 2013; Paxinos & Watson, 2007). Labeling in these regions was sparse in rostral cases and absent in caudal cases. No cases contained labeling in the dorsal lateral geniculate nucleus. Immediately ventral to the medial geniculate nucleus, we found moderately dense labeling in the peripeduncular nucleus.

### Midbrain

In rostral cases, the tectum contained one of the densest targets in the brain. There, we found a thin, dense stripe of Syp-mCherry-labeled boutons near the midline of the deep layers of the superior and inferior colliculi. This terminal field, immediately dorsal to the PAG, was apparent in serial coronal sections (**Figure 11a-b**) and even more prominent in the sagittal plane (**Figure 11c-d**). This rostrocaudally elonged zone appears to be the “tectal longitudinal column” (TLC) described in rats (Saldana et al., 2007). While the TLC was one of the most prominent targets in the brain in rostral cases, we found virtually no labeling in this region in caudal cases. Labeled axons and boutons reached this target zone via a periventricular pathway, through the periaqueductal gray matter, which contained moderately dense labeling as well.

**Figure 11.**
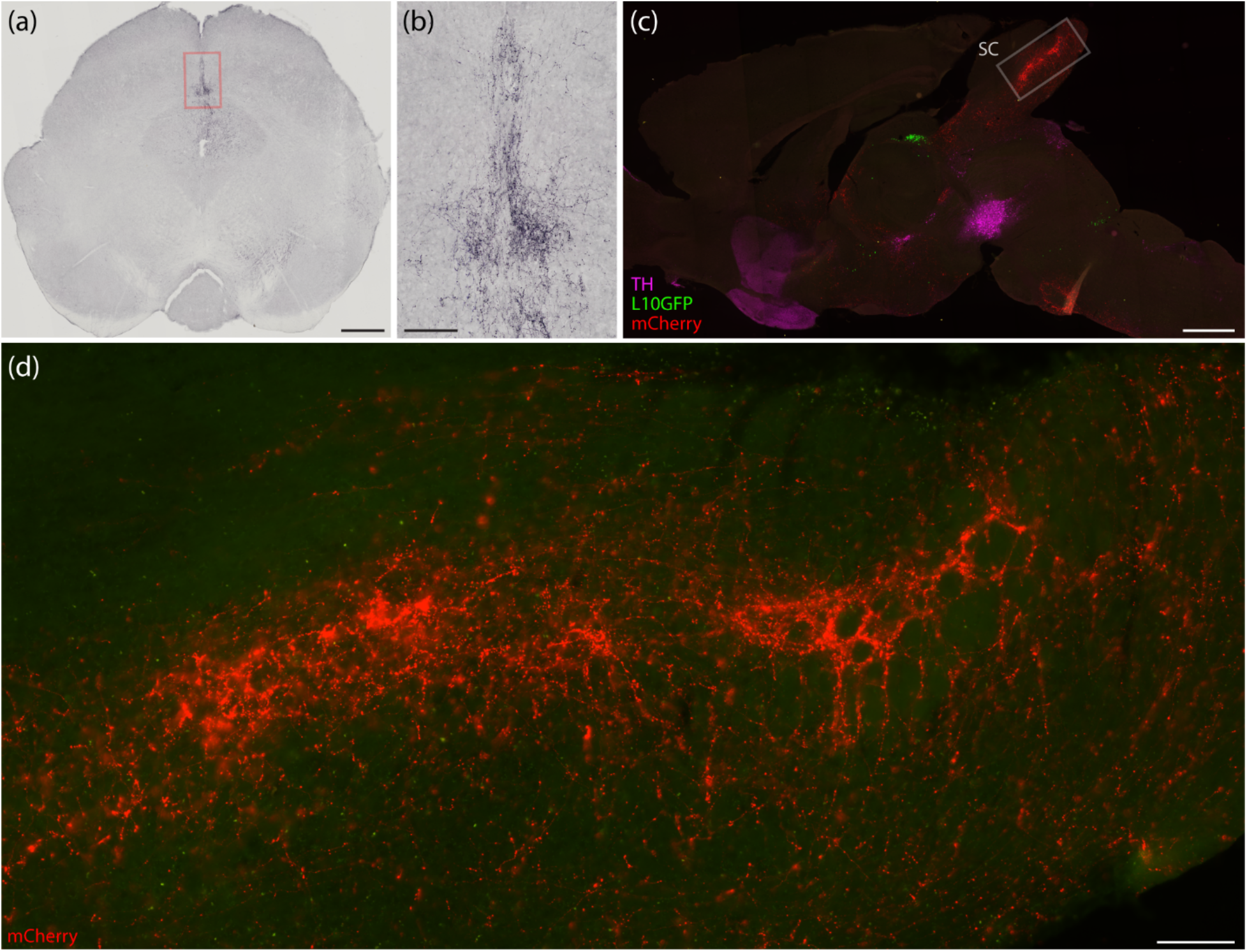
(a–b) Dense labeling in the TLC (case 5569; coronal section, NiDAB labeling for Syp-mCherry). (c) Sagittal view of median collicular labeling with L10GFP Cre-reporter for *Nps* (green) plus immunofluorescence labeling for TH (magenta; case 6731) and magnified area of interest in (d). For the original description of this region in rats, see Figures 2–3 of (Saldana et al., 2007). Scalebars are (a) 500 µm, (b) 100 µm, (c) 1 mm, and (d) 100 µm.

Separate from this periventricular pathway, the midbrain received moderate labeling along a ventrolateral pathway. Most axons coursed alongside the medial lemniscus and dorsal to the substantia nigra without arborizing in the midbrain, but many produced a concentration of branches and boutons in the retrorubral field and extended rostrally into the peripeduncular and subparafasciular thalamic target zones described above.

### Cerebellum

No part of the cerebellar cortex or deep cerebellar nuclei contained labeling in any cases.

### Hindbrain

While NPS neurons supply a descending pathway, these projections did not produce wide-ranging axonal labeling in the hindbrain. A thin rim of moderately dense labeling encased the superior olivary nucleus, trapezoid bodies, and ventral nucleus of the lateral lemniscus. We also found light labeling in the ventrolateral pontine reticular nucleus and forming a rim around the facial motor nucleus. Moderately dense labeling also targeted the superior salivatory nucleus, just rostral and dorsolateral to the facial motor nucleus. The nucleus of the solitary tract had no more than trace labeling. A region of the caudal ventrolateral medulla that included the A1 group and caudal pressor area (CPA) contained sparse labeling.

## Discussion

Glutamatergic neurons in the PB form two macropopulations, an *Atoh1*-derived macropopulation and an *Lmx1*-expressing macropopulation. PB neurons project to target regions via four pathways: a periventricular pathway, the central tegmental tract, a ventral pathway, and a descending pathway (**Figure 12a**). Neurons in the *Lmx1* macropopulation project axons primarily via the central tegmental tract, while neurons in the *Atoh1*-derived macropopulation project axons primarily via the ventral pathway. NPS neurons are a subset of the *Atoh1* macropopulation and project axons via this same ventral pathway, as well as a robust periventricular pathway (**Figure 12b**). We found that *Nps*-expressing PB neurons project heavily to the midline thalamus, medial hypothalamus, and midline tectum. In contrast, we found little to no labeling in the cerebral cortex, olfactory bulb, amygdala, and cerebellum. After discussing limitations of our study, we compare our results to previous work and forecast the function of these neurons.

**Figure 12.**
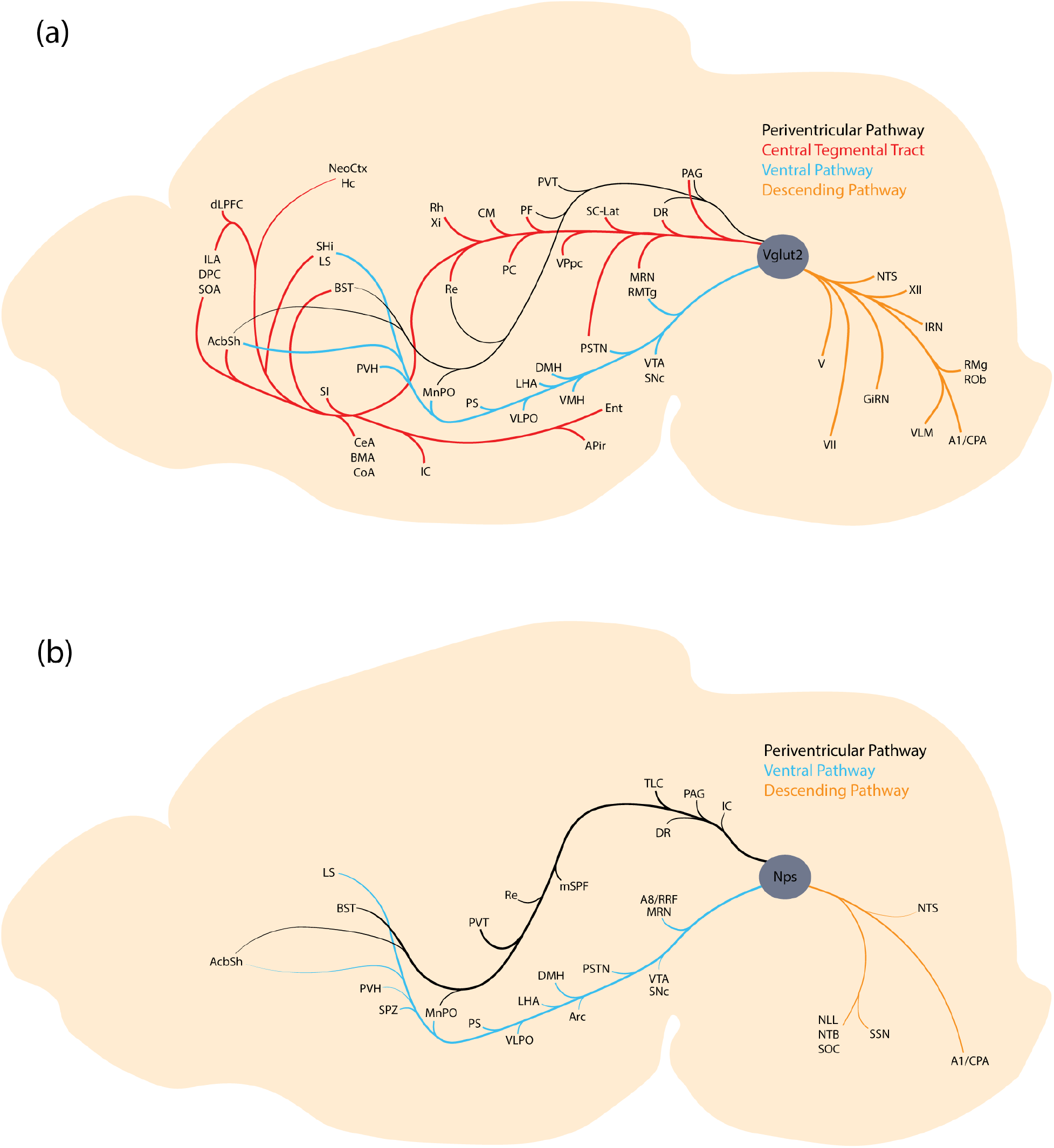
Projection pathways and targets of (a) PB glutamatergic neurons in comparison to (b) the *Nps*-expressing subset of glutamatergic neurons. *Nps*-expressing neurons project prominently via a periventricular pathway.

### Limitations

Syp-mCherry tracing with BoutonNet algorithmic processing has several advantages compared to previous anterograde tracing techniques (Grady et al., 2022). Syp-mCherry concentrates in the presynaptic bouton, while previous tracers uniformly distribute in the axoplasm. However, a small portion of the putative boutons labeled by our method likely represent non-synaptic portions of axons cut in cross-section because Syp-mCherry must travel through the axon shaft to reach presynaptic boutons. To mitigate this limitation, we also visually inspected Syp-mCherry labeling in every brain region and applied semi-quantitative density standards used in previous tracing studies (Huang et al., 2020; Huang et al., 2021). Despite these efforts to identify boutons from the morphology of labeling, determining whether individual boutons form structural synapses would require the use of electron microscopy (Wouterlood & Jorritsma-Byham, 1993), and determining whether they form functional synapses would require the use of electrophysiology (Petreanu et al., 2007). Neither of these approaches is practical at the scale of this study.

We genetically targeted NPS neurons using a knockin-Cre strain with a 2A “self-cleaving” peptide inserted after the last exon of the endogenous *Nps* gene (Huang et al., 2022). We used a Cre-conditional construct (DIO-Syp-mCherry), so only neurons with sufficient Cre expression between the time of injection and perfusion were transduced. We performed these experiments in adult mice, so the results reflect axonal projections of neurons with adult *Nps* expression, not just developmental expression, which may be more widespread.

Additionally, the Syp-mCherry promoter is different from the *Nps* gene promoter. Therefore, neurons with sufficient Cre expression to activate the DIO construct produce large amounts of Syp-mCherry, without reflecting varying levels of *Nps* expression in those neurons. This may be relevant for NPS neurons due to the known variations in NPS protein and *Nps* mRNA expression with physiological state. Thus, while our approach has high sensitivity for neurons that express *Nps*, the axonal labeling density in our results does not reflect *Nps* expression level in those neurons.

Although we compared the projection patterns of the rostral and caudal group of NPS neurons, we did not distinguish subpopulations of neurons within these groups. The rostral group of NPS neurons concentrates between the tip of the superior cerebellar peduncle and the medial edge of the lateral lemniscus, resembling the cytoarchitecturally defined “extreme lateral” subnucleus in the rat PB (Fulwiler & Saper, 1984). However, this group also includes neurons scattered caudally through the lateral PB, and additional neurons expressing an *Nps* Cre-reporter extend rostrally along the medial edge of the lateral lemniscus, dorsal to the population of TIP39/*Pth2*-expressing neurons in the medial paralemniscal nucleus (Dobolyi et al., 2003; Sun et al., 2022; Varga et al., 2008) and overlapping a location described as the “semilunar nucleus” in rats (Gómez-Martínez et al., 2023). Neurons concentrated in the main (“extreme lateral”) NPS cluster express high levels of *Nps* mRNA, while the other neurons identifiable by Cre-reporter expression contain very little, so these neurons may form separate subpopulations with separate connections and functions. Discriminating differences in projection patterns would require first identifying additional genetic markers that can distinguish NPS subpopulations. Within the caudal cluster, most NPS neurons near the locus coeruleus express *Foxp2*, but Barrington’s nucleus includes a small number of neurons with *Nps* Cre-reporter expression that lack FoxP2 (Huang et al., 2022). Differentiating the projection pattern of Barrington’s NPS neurons that lack FoxP2 from that of peri-LC NPS neurons that express *Foxp2* may be possible with intersectional genetic targeting.

### Comparison with previous neuroanatomical literature

Generally, the pattern of Syp-mCherry labeling from NPS neurons in the PB region approximated the pattern of NPS fiber immunolabeling (**Figure 2**), as described in a previous study using the same NPS antiserum (Clark et al., 2011). This implies that neurons in the PB region supply most of the NPS peptide in the mouse brain. Neighboring, *Cck*-expressing neurons in the PB project heavily to the VMH (Garfield et al., 2014; Grady et al., 2020), and a previous study of NPS immunoreactivity reported dense fiber labeling in this hypothalamic nucleus (Clark et al., 2011). However, the VMH contained virtually no NPS-immunoreactive fibers and was largely devoid of Syp-mCherry axonal labeling in our study (**Figure 2e**).

Additionally, we identified output projections to regions of the brain that contain sparse immunolabeling, including the ventral BST, mSPF, and TLC. These target regions contained dense Syp-mCherry labeling and appear to be important targets for NPS neurons. A previous study in rats observed retrograde labeling in the “extreme lateral” PB subnucleus after a tracer injection spanning the rostral BST, nucleus accumbens, and lateral septum (Fulwiler & Saper, 1984). Another previous tracing study identified retrograde labeling in the PB region after tracer injections into the mSPF (Wang et al., 2006), but we are not aware of any previous retrograde tracing studies that described the dense connectivity from the PB region to TLC. This reigon has *Npsr1* expression as part of a broader pattern of expression in the superior colliculi (Clark et al., 2011). To the best of our knowledge, the presence or absence *Npsr1* expression in the mSPF has not been clarified in previous work.

NPS neurons are glutamatergic, *Atoh1*-derived, and *Foxp2*-expressing, but they do not express *Pdyn*. (Huang et al., 2022; Xu et al., 2007). The present results are largely consistent with previous tracing data on the efferent projections of *Foxp2*-expressing neurons in rats and mice, including a general lack of labeling in the olfactory bulb, cerebral cortex, amygdala (Huang et al., 2020; Shin et al., 2011). Our results contrast the projections of the PB *Lmx1* macropopulation, including *Calca*-expressing neurons, which primarily target the amygdala and cerebral cortex and send much less output to the hypothalamus (Huang et al., 2021).

Most NPS projections identified here represent a subset of the efferent projections previously identified in our study of the larger population of glutamatergic *Foxp2*-expressing neurons in the PB region (Huang et al., 2020). However, NPS neurons densely targeted a small number of brain regions, including the TLC and NLL, that were less evident in our previous study. Interestingly, we found dense fiber labeling in the TLC in Atoh1-Cre mice with a tdTomato Cre-reporter (see Figure 20b of Karthik et al., 2022). The injection sites in our *Foxp2* tracing study were centered caudal to the main concentration of NPS neurons, and the presence of labeling in the TLC in Atoh1-Cre tdTomato reporter mice supports the present finding that *Atoh1*-derived, *Nps*-expressing neurons in the rostral PB project to this region.

The median superior colliculus contains the tectal longitudinal column with ventral (TLCv) and dorsal (TLCd) components (Aparicio & Saldaña, 2014). Anterograde Syp-mCherry axonal labeling from *Nps-*expressing neurons in the PB appear to target TLCv with less dense labeling dorsally in the TLCd (**Figure 11**). Interestingly, the periolivary region, which receives input from the NPS neurons, appears to be reciprocally connected with the TLCv (Saldaña et al., 2009; Viñuela et al., 2011).

### Functional implications

The PB is a major integrator of interoceptive sensory inputs. The genetic identity and location of neurons in the PB is a useful way to delineate their different projections and functions. For example, *Pdyn*-expressing neurons in the dorsolateral PB relay warm thermosensory information (Geerling et al., 2016), Satb2-expressing neurons in the medial PB relay taste information (Fu et al., 2019; Jarvie et al., 2021), and *Calca*-expressing neurons in the external lateral subdivision induce taste aversion and inhibit appetite (Carter et al., 2015; Carter et al., 2013; Palmiter, 2018). In contrast to these known populations, *Nps*-expressing neurons are a neuroanatomically distinct group with a novel projection pattern. They target brain regions that regulate circadian rhythm and threat response.

The SCN is the master circadian regulator, and among brain regions with direct connectivity to the SCN, NPS axons project to the SPZ, PVT, DMH, and IGL. The SPZ is the main output target of the SCN and plays an important role in the circadian regulation of locomotor activity, body temperature, and sleep (Vujovic et al., 2015). Evidence exists for a relationship between NPS and sleep: a human *NPSR1* gain-of-function mutation was associated with substantially reduced sleep, and mice carrying this mutation also slept less (Xing et al., 2019). Additionally, central injection of an NPSR1 antagonist reduced wakefulness in rats (Oishi et al., 2014). Another region of dense labeling in the midline thalamus was the mSPF. In our analysis of NPS neuron anterograde tracing, we were surprised to find dense labeling in the mSPF, whose neurons are medial and ventral to the fasciculus retroflexus and have not been comprehensively studied. This compact and unusual thalamic nucleus contains GABAergic neurons and sends a prominent output projection to the IGL (Vrang et al., 2003), which is implicated in circadian function due to its input from the retina and output to the SCN (Moore et al., 2000). In addition to the mSPF, the IGL also receives light axonal input from the NPS neurons. The anterior PVT and DMH also play important roles in circadian regulation, and each of these sites receives heavy input from NPS neurons. It is likely that NPS projections to these brain regions influence circadian patterns.

Adaptive responses to threats are fundamental to survival. NPS axons target several brain regions that are implicated in modulating threat responses: the DMH, ventral BST, PVT, and medial tectum. NPS axons blanked the medial hypothalamus while largely avoiding the VMH, mamillary bodies, and SCN, and they produced particularly dense labeling in the DMH. *Cck*-expressing neurons in the superior lateral PB project to the VMH and are activated by noxious, aversive input from the spinal cord (Hermanson et al., 1998), and neurons in the VMH cause aggressive attack behavior in male mice (Lin et al., 2011). However, the lack of Syp-mCherry labeling in the VMH and the mutual exclusivity of *Cck* and *Nps* expression suggest that NPS neurons do not regulate these behaviors. Dense labeling in the DMH spreads dorsally and laterally into the zona incerta and encompasses the A13 catecholaminergic cell group. Electrically stimulating this medial region of the hypothalamus triggers escape behaviors, while stimulating the lateral hypothalamus triggers exploratory locomotion (Lammers et al., 1988). The DMH contains *Npsr1* expression, and central NPS injections increase locomotion, so it is tempting to speculate that NPS-DMH connectivity is responsible for escape responses to threats.

NPS axons avoided the dorsal BST, while broadly targeting a subregion of the ventral BST that receives AgRP input from the arcuate hypothalamic nucleus. The BST appears to play a role in sustained responses to threat (Davis et al., 2010; Gewirtz et al., 1998; Walker et al., 2003) and is implicated in general anxiety disorder in humans (Lebow & Chen, 2016; Yassa et al., 2012). Based on associations between human *NPSR1* gain-of-function mutations with panic and anxiety disorders (Domschke et al., 2011; Donner et al., 2010; Okamura et al., 2007; Pape et al., 2010), NPS projections to the BST may play a role in regulating long-term responses to threats.

The PVT is the most prominent region with Syp-mCherry labeling, as well as NPS-immunoreactive fibers and *Npsr1* mRNA expression (Clark et al., 2011). Notably, activating PB projections to the PVT is aversive (Zhu et al., 2022). Also, deleting either *Nps* or *Npsr1* attenuates conditioned fear responses, and these responses can be rescued by injecting NPS into the PVT (Garau et al., 2022). NPS neuron connectivity to the PVT is probably important for relaying aversive information.

In the midbrain, we found a dense patch of labeling in the midline of the tectum. This labeling was most prominent in cases with rostral injections sites and virtually absent in caudal cases. This area of the brain was identified in rats as the TLC (Saldana et al., 2007). Neurons in this region may be important for encoding threat salience, as opposed neurons in the dorsal PAG that regulate escape behavior (Evans et al., 2018). We postulate that NPS axons targeting the medial tectum are part of a neural circuit that encodes for threat salience.

Descending projections to the hindbrain targeted the superior salivatory nucleus, while skirting around the edges of the VII. Via their peripheral output projections to the pterygopalatine ganglion, SSN neurons control the lacrimal gland, along with mucous secretion and dilation of cerebral blood vessels. Along with their output to other target regions implicated in aversive responses, this connection to the SSN may allow NPS neurons to drive autonomic responses to distress, such as mucous secretion from the nasal mucosa and tear secretion from the lacrimal glands.

### Conclusion

In pharmacological studies, NPS promotes arousal and increases locomotion.

Receptor mutations are associated with reduced sleep time, as well as panic and anxiety disorders. In this study, we mapped and analyzed the output projections of NPS neurons in the PB region. Their target regions predict that NPS neurons influence circadian rhythms and compute responses to threat.

### Role of authors

JCG planned the project and secured funding. SG bred, weaned, and genotyped Nps-2A-Cre mice and Nps-2A-Cre;R26-lsl-L10GFP mice. RZ and SG performed histologic staining and slide microscopy. SG performed stereotaxic injections. RZ and DH drafted the figures. RZ, DH, and JCG edited the figures and figure legends. RZ and JCG drafted and edited the text of the paper.

### Data Availability

The data that support the findings of this study are available from the corresponding author upon reasonable request.

### Other Acknowledgements

This work was supported by an Accelerator Award from the Iowa Neuroscience Institute. Funding support for Richie Zhang was provided in 2022 by a Summer Research Fellowship award from the Carver College of Medicine and in 2023 by the National Heart, Lung, and Blood Institute of the NIH under award number 1T35HL166206. Knockin Nps-2A-Cre mice were generated at the University of Iowa Genome Editing Core Facility, which is supported in part by grants from the NIH and from the Roy J. and Lucille A. Carver College of Medicine – we thank Bill Paradee, Norma Sinclair, Patricia Yarolem, Joanne Schwarting, and Rongbin Guan for their technical expertise in generating these mice. Finally, we thank Fillan S. Grady for help with implementing the BoutonNet program.

### Conflict of Interest

The authors declare no conflicts of interest.

## Supporting information

Supplemental Figure 1 (full rostrocaudal injection sites, Syp-mCherry NiDAB labeling with Nissl counterstain)

Supplemental video (CCFv3 mouse brain distribution of neurons with a history of Nps expression)

## Abbreviations

A1: = A1 catecholaminergic group
A8: = A8 catecholaminergic group
ac: = anterior commissure
AcbSh: = accumbens nucleus, shell
ACA: = anterior cingulate cor7cal area
AgRP: = agou7-related pep7de
AHA: = anterior hypothalamic area
AI: = agranular insular cortex
AmbC: = ambiguus nucleus compact part
AON: = anterior olfactory nucleus
AP: = area postrema
APir: = amygdalopiriform transi7on area
APN: = anterior pretectal nucleus
Arc: = arcuate nucleus
AVP: = arginine vasopressin
BLA: = basolateral amygdalar nucleus
BMA: = basomedial amygdalar nucleus
BST: = bed nucleus of the stria terminalis
BSTaL: = BST anterolateral subnucleus
BSTam: = BST anteromedial subnucleus
BSTfu: = BST fusiform subnucleus
BSTov: = BST oval subnucleus
BSTp: = BST posterior subnucleus
CeA: = central nucleus of the amygdala
CeAc: = CeA capsular subdivision
CeAl: = CeA lateral subdivision
CeAm: = CeA medial subdivision
CGRP: = calcitonin gene-related pep7de
Cla: = claustrum
CLi: = central linear raphe
CM: = central medial thalamic nucleus
CoA: = cor7cal amygdalar nucleus
CPA: = caudal pressor area
Cun: = cuneiform nucleus
DCN: = deep cerebellar nuclei
DCO: = dorsal cochlear nucleus
Dl: = dysgranular insular cortex
DMH: = dorsomedial hypothalamic nucl.
DP: = dorsal peduncular cor7cal area
DR: = dorsal raphe nucleus
DTN: = dorsal tegmental nucleus
Ent =: entorhinal cor7cal area
Epd/Epv: = endopiriform nucleus, dorsal/ventral subdivisions
Epy: = ependymal cell layer of cerebral aqueduct
EW: = Edinger-Westphal nucleus
FL: = flocculus of the cerebellum
fr: = fasciculus retroflexus
fx: = fornix
GI =: granular insular cortex
GiRN: = gigantocellular re7cular nucleus
GPe: = globus pallidus external
GPi: = globus pallidus internal
GRP: = gastrin-related pep7de
Hc: = hippocampus
IA: = intercalated amygdalar nucleus
IC: = inferior colliculus
IFV: = interfascicular trigeminal nucleus
IG: = induseum griseum
IGL: = intergeniculate leaflet
ILA: = infralimbic cor7cal area
IMD: = intermediodorsal thalamic nucl.
IO: = inferior olivary complex
IPN: = interpeduncular nucleus
IRN: = intermediate re7cular nucleus
L6 PFC: = layer 6 prefrontal cortex
LAr: = lateral amygdaloid nucleus
LDT: = laterodorsal tegmental nucleus
LGd: = lateral geniculate nucleus, dorsal subdivision
LGv: = lateral geniculate nucleus, ventral subdivision
LHA: = lateral hypothalamic area
LHb: = lateral habenular nucleus
LM: = lateral mammillary nucleus
LPOA: = lateral preop7c area
LRN: = lateral re7cular nucleus
LSi: = lateral septum, intermediate subdivision
LSr: = lateral septum, rostral subdivision
LSv: = lateral septum ventral subdivision
MA: = magnocellular nucleus
MB: = mammillary bodies
MD: = mediodorsal thalamic nucleus
MeA: = medial amgydalar nucleus
MGN: = medial geniculate nucleus
MHb: = medial habenular nucleus
MM: = medial mammillary nucleus
MnPO: = median preop7c nucleus
MOB: = main olfactory bulb
MPN: = medial preop7c nucleus
mPOA: = medial preop7c area
MR: = median raphe
MRF: = midbrain re7cular forma7on
MS: = medial septum
mSPF: = magnocellular subparafascicular nucleus
NBM: = nucleus basalis of Meynert
NDB: = nucleus of the diagonal band
NeoCtx: = neocortex
NLL: = nucleus of the lateral lemniscus
NPS: = neuropep7de S
NTB: = nucleus of trapezoid body
NTS: = nucleus of the solitary tract
NTS-Lat: = NTS lateral subdivision
NTS-Med: = NTS medial subdivision
OVLT: = organum vasculosum of lamina terminalis
PAG: = Periaqueductal gray maMer
PAG-L: = PAG lateral subdivision
PAGdl: = PAG dorsolateral subdivision
PAGdm: = PAG dorsomedial subdivision
PAGvL: = PAG ventrolateral subdivision
PaRN: = parvicellular re7cular nucleus
PB: = parabrachial nucleus
PC: = paracentral thalamic nucleus
PeriRh =: perirhinal cor7cal area
PF: = parafascicular thalamic nucleus
PG: = pon7ne grey
PH: = posterior hypothalamic nucleus
Pir: = piriform cor7cal area
PL: = prelimbic cor7cal area
PLi: = posterior limitans thalamic nucleus
PMd/v: = posterior mammillary nucleus, dorsal/ventral
Po: = posterior thalamic complex
PP: = peripeduncular nucleus
PPN: = pedunculopon7ne tegmental nucleus
PreT: = pretectal region
PRN: = pon7ne re7cular nucleus
PS: = parastrial nucleus
PSTN: = parasubthalamic nucleus
PSV: = principal sensory trigeminal nucl.
PT: = paratenial thalamic nucleus
PVH: = paraventricular hypothalamic nucleus
PVT: = paraventricular thalamic nucleus
RCA: = retrochiasma7c area
Re: = reuniens thalamic nucleus
Rh: = rhomboid thalamic nucleus
RLi: = rostral linear raphe
RMg/Ob/Pa: = raphe magnus/obscurus/ pallidus
RMTg: = rostromedial tegmental nucleus
RN: = red nucleus
RR: = retrorubral area
SC: = superior colliculus
SC-Lat: = SC lateral aspect
SC-Sen: = SC sensory-related
SCN: = suprachiasma7c nucleus
SCO: = subcommissural organ
SEZ: = subependymal germinal zone
SF: = septofimbral nucleus
SFO: = subfornical organ
SHi: = septohippocampal nucleus
SI: = Subs7an7a inominata
SNc: = Subs7an7a nigra pars compacta
SNr: = Subs7an7a nigra pars re7cularis
SOC: = superior olivary complex
SON: = supraop7c nucleus
SPFp: = subparafasicular nucleus, parvicellular part
SpV: = spinal trigeminal nucleus
SPZ: = subparaventricular zone
SSN: = superior salivatory nucleus
STN: = subthalamic nucleus
Str (CP): = striatum (caudoputamen)
Str/GP bdr: = striatum/pallidum border
SUM: = supramammillary nucleus
TH: = tyrosine hydroxylase
TLC: = tectal longitudinal column
TM: = tuberomammillary nucleus
TRN: = tegmental re7cular nucleus
TR: = amygdalopiriform transi7on area
TT: = tenia tecta
V: = trigeminal motor nucleus
VII: = facial motor nucleus
VLPO: = ventrolateral preop7c nucleus
VM: = ventromedial thalamic nucleus
VMH: = ventromedial hypothalamic nucl.
VNC: = ves7bular nuclear complex
VPpc: = parvicellular ventral posterior thalamic nucleus
VTA: = ventral tegmental area
Xi: = xiphoid thalamic nucleus
XII: = hypoglossal motor nucleus
ZI: = zona incer

---

**Supplementary Video 1.** Rotations of a 3D model of NPS neuron locations in the mouse brain.

**Supplemental Figure 1.**
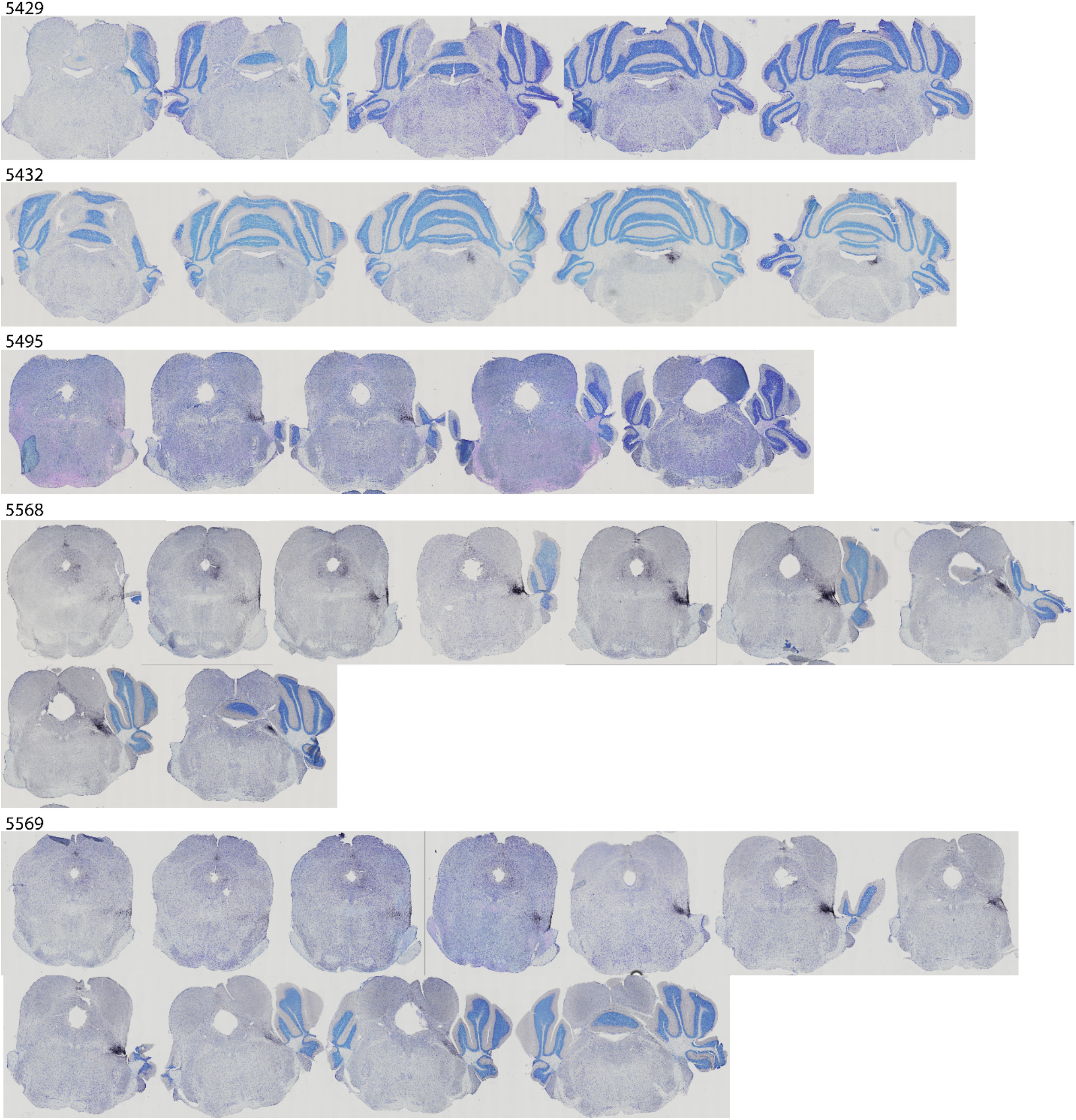
Full rostral-to-caudal Nissl-counterstained sections with Syp-mCherry NiDAB immunolabeling in all injection sites cut in the coronal plane.

## Notes

### Competing Interest Statement

The authors have declared no competing interest.

## References

Adori, C., Barde, S., Bogdanovic, N., Uhlen, M., Reinscheid, R. R., Kovacs, G. G., & Hokfelt, T. (2015). Neuropeptide S- and Neuropeptide S receptor-expressing neuron populations in the human pons. Front Neuroanat, 9, 126. https://doi.org/10.3389/fnana.2015.00126

Adori, C., Barde, S., Vas, S., Ebner, K., Su, J., Svensson, C., . . . Hökfelt, T. (2016). Exploring the role of neuropeptide S in the regulation of arousal: a functional anatomical study. Brain Struct Funct, 221(7), 25. https://doi.org/10.1007/s00429-015-1117-5

Aparicio, M. A., & Saldaña, E. (2014). The dorsal tectal longitudinal column (TLCd): a second longitudinal column in the paramedian region of the midbrain tectum. Brain Struct Funct, 219(2), 607–630. https://doi.org/10.1007/s00429-013-0522-x

Carter, M. E., Han, S., & Palmiter, R. D. (2015). Parabrachial calcitonin gene-related peptide neurons mediate conditioned taste aversion. J Neurosci, 35(11), 4582–4586. https://doi.org/10.1523/JNEUROSCI.3729-14.2015

Carter, M. E., Soden, M. E., Zweifel, L. S., & Palmiter, R. D. (2013). Genetic identification of a neural circuit that suppresses appetite. Nature, 503(7474), 111–114. https://doi.org/10.1038/nature12596

Cechetto, D. F., Standaert, D. G., & Saper, C. B. (1985). Spinal and trigeminal dorsal horn projections to the parabrachial nucleus in the rat. J Comp Neurol, 240(2), 153–160. https://doi.org/10.1002/cne.902400205

Clark, S. D., Duangdao, D. M., Schulz, S., Zhang, L., Liu, X., Xu, Y. L., & Reinscheid, R. K. (2011). Anatomical characterization of the neuropeptide S system in the mouse brain by in situ hybridization and immunohistochemistry. J Comp Neurol, 519(10), 1867–1893. https://doi.org/10.1002/cne.22606

Davis, M., Walker, D. L., Miles, L., & Grillon, C. (2010). Phasic vs sustained fear in rats and humans: role of the extended amygdala in fear vs anxiety. Neuropsychopharmacology, 35(1), 105–135. https://doi.org/10.1038/npp.2009.109

Dobolyi, A., Palkovits, M., & Usdin, T. B. (2003). Expression and distribution of tuberoinfundibular peptide of 39 residues in the rat central nervous system. J Comp Neurol, 455(4), 547–566. https://doi.org/10.1002/cne.10515

Domschke, K., Reif, A., Weber, H., Richter, J., Hohoff, C., Ohrmann, P., . . . Deckert, J. (2011). Neuropeptide S receptor gene -- converging evidence for a role in panic disorder. Mol Psychiatry, 16(9), 938–948. https://doi.org/10.1038/mp.2010.81

Dong, H. W. (2008). Allen reference atlas : a digital color brain atlas of the C57Black/6J male mouse. Wiley.

Donner, J., Haapakoski, R., Ezer, S., Melen, E., Pirkola, S., Gratacos, M., . . . Hovatta, I. (2010). Assessment of the neuropeptide S system in anxiety disorders. Biol Psychiatry, 68(5), 474–483. https://doi.org/10.1016/j.biopsych.2010.05.039

Duangdao, D. M., Clark, S. D., Okamura, N., & Reinscheid, R. K. (2009). Behavioral phenotyping of neuropeptide S receptor knockout mice. Behav Brain Res, 205(1), 1–9. https://doi.org/10.1016/j.bbr.2009.07.024

Ensho, T., Nakahara, K., Suzuki, Y., & Murakami, N. (2017). Neuropeptide S increases motor activity and thermogenesis in the rat through sympathetic activation. Neuropeptides, 65, 21–27. https://doi.org/10.1016/j.npep.2017.04.005

Evans, D. A., Stempel, A. V., Vale, R., Ruehle, S., Lefler, Y., & Branco, T. (2018). A synaptic threshold mechanism for computing escape decisions. Nature, 558(7711), 590–594. https://doi.org/10.1038/s41586-018-0244-6

Fendt, M., Buchi, M., Bürki, H., Imobersteg, S., Ricoux, B., Suply, T., & Sailer, A. W. (2011). Neuropeptide S receptor deficiency modulates spontaneous locomotor activity and the acoustic startle response. Behav Brain Res, 217(1), 1–9. https://doi.org/10.1016/j.bbr.2010.09.022

Fu, O., Iwai, Y., Kondoh, K., Misaka, T., Minokoshi, Y., & Nakajima, K. I. (2019). SatB2-Expressing Neurons in the Parabrachial Nucleus Encode Sweet Taste. Cell Rep, 27(6), 1650–1656 e1654. https://doi.org/10.1016/j.celrep.2019.04.040

Fulwiler, C. E., & Saper, C. B. (1984). Subnuclear organization of the efferent connections of the parabrachial nucleus in the rat. Brain Res, 319(3), 229–259.

Garau, C., Liu, X., Calo, G., Schulz, S., & Reinscheid, R. K. (2022). Neuropeptide S Encodes Stimulus Salience in the Paraventricular Thalamus. Neuroscience, 496, 83–95. https://doi.org/10.1016/j.neuroscience.2022.06.013

Garfield, A. S., Shah, B. P., Madara, J. C., Burke, L. K., Patterson, C. M., Flak, J., . . . Heisler, L. K. (2014). A parabrachial-hypothalamic cholecystokinin neurocircuit controls counterregulatory responses to hypoglycemia. Cell Metab, 20(6), 1030–1037. https://doi.org/10.1016/j.cmet.2014.11.006

Gasparini, S., Resch, J. M., Narayan, S. V., Peltekian, L., Iverson, G. N., Karthik, S., & Geerling, J. C. (2019). Aldosterone-sensitive HSD2 neurons in mice. Brain Struct Funct, 224(1), 387–417. https://doi.org/10.1007/s00429-018-1778-y

Geerling, J. C., Kim, M., Mahoney, C. E., Abbott, S. B., Agostinelli, L. J., Garfield, A. S., . . . Scammell, T. E. (2016). Genetic identity of thermosensory relay neurons in the lateral parabrachial nucleus. Am J Physiol Regul Integr Comp Physiol, 310(1), R41–54. https://doi.org/10.1152/ajpregu.00094.2015

Gewirtz, J. C., McNish, K. A., & Davis, M. (1998). Lesions of the bed nucleus of the stria terminalis block sensitization of the acoustic startle reflex produced by repeated stress, but not fear-potentiated startle. Prog Neuropsychopharmacol Biol Psychiatry, 22(4), 625–648. https://doi.org/10.1016/s0278-5846(98)00028-1

Gottlieb, D. J., O’Connor, G. T., & Wilk, J. B. (2007). Genome-wide association of sleep and circadian phenotypes. BMC Med Genet, 8 *Suppl 1*, S9. https://doi.org/10.1186/1471-2350-8-S1-S9

Grady, F., Peltekian, L., Iverson, G., & Geerling, J. C. (2020). Direct Parabrachial-Cortical Connectivity. Cereb Cortex. https://doi.org/10.1093/cercor/bhaa072

Grady, F. S., Graff, S. A., Aldridge, G. M., & Geerling, J. C. (2022). BoutonNet: an automatic method to detect anterogradely labeled presynaptic boutons in brain tissue sections. Brain Struct Funct. https://doi.org/10.1007/s00429-022-02504-y

Gómez-Martínez, M., Rincón, H., Gómez-Álvarez, M., Gómez-Nieto, R., & Saldaña, E. (2023). The nuclei of the lateral lemniscus: unexpected players in the descending auditory pathway [Original Research]. Frontiers in Neuroanatomy, 17. https://doi.org/10.3389/fnana.2023.1242245

Herbert, H., Moga, M. M., & Saper, C. B. (1990). Connections of the parabrachial nucleus with the nucleus of the solitary tract and the medullary reticular formation in the rat. J Comp Neurol, 293(4), 540–580.

Hermanson, O., Larhammar, D., & Blomqvist, A. (1998). Preprocholecystokinin mRNA-expressing neurons in the rat parabrachial nucleus: subnuclear localization, efferent projection, and expression of nociceptive-related intracellular signaling substances. J Comp Neurol, 400(2), 255–270.

Huang, D., Grady, F. S., Peltekian, L., & Geerling, J. C. (2020). Efferent Projections of Vglut2, Foxp2 and Pdyn Parabrachial Neurons in Mice. J Comp Neurol. https://doi.org/10.1002/cne.24975

Huang, D., Grady, F. S., Peltekian, L., Laing, J. J., & Geerling, J. C. (2021). Efferent projections of CGRP/Calca-expressing parabrachial neurons in mice. J Comp Neurol. https://doi.org/10.1002/cne.25136

Huang, D., Zhang, R., Gasparini, S., McDonough, M. C., Paradee, W. J., & Geerling, J. C. (2022). Neuropeptide S (NPS) neurons: Parabrachial identity and novel distributions. J Comp Neurol. https://doi.org/10.1002/cne.25400

Jarvie, B. C., Chen, J. Y., King, H. O., & Palmiter, R. D. (2021). Satb2 neurons in the parabrachial nucleus mediate taste perception. Nat Commun, 12(1), 224. https://doi.org/10.1038/s41467-020-20100-8

Karthik, S., Huang, D., Delgado, Y., Laing, J. J., Peltekian, L., Iverson, G. N., . . . Geerling, J. C. (2022). Molecular ontology of the parabrachial nucleus. J Comp Neurol. https://doi.org/doi.org/10.1002/cne.25307

Laitinen, T., Polvi, A., Rydman, P., Vendelin, J., Pulkkinen, V., Salmikangas, P., . . . Kere, J. (2004). Characterization of a common susceptibility locus for asthma-related traits. Science, 304(5668), 300–304. https://doi.org/10.1126/science.1090010

Lammers, J. H., Kruk, M. R., Meelis, W., & van der Poel, A. M. (1988). Hypothalamic substrates for brain stimulation-induced patterns of locomotion and escape jumps in the rat. Brain Res, 449(1-2), 294–310. https://doi.org/10.1016/0006-8993(88)91045-1

Lebow, M. A., & Chen, A. (2016). Overshadowed by the amygdala: the bed nucleus of the stria terminalis emerges as key to psychiatric disorders. Mol Psychiatry, 21(4), 450–463. https://doi.org/10.1038/mp.2016.1

Leonard, S. K., Dwyer, J. M., Sukoff Rizzo, S. J., Platt, B., Logue, S. F., Neal, S. J., . . . Ring, R. H. (2008). Pharmacology of neuropeptide S in mice: therapeutic relevance to anxiety disorders. Psychopharmacology (Berl), 197(4), 601–611. https://doi.org/10.1007/s00213-008-1080-4

Li, W., Chang, M., Peng, Y. L., Gao, Y. H., Zhang, J. N., Han, R. W., & Wang, R. (2009). Neuropeptide S produces antinociceptive effects at the supraspinal level in mice. Regul Pept, 156(1-3), 90–95. https://doi.org/10.1016/j.regpep.2009.03.013

Lin, D., Boyle, M. P., Dollar, P., Lee, H., Lein, E. S., Perona, P., & Anderson, D. J. (2011). Functional identification of an aggression locus in the mouse hypothalamus. Nature, 470(7333), 221–226. https://doi.org/10.1038/nature09736

Liu, X., Zeng, J., Zhou, A., Theodorsson, E., Fahrenkrug, J., & Reinscheid, R. K. (2011). Molecular fingerprint of neuropeptide S-producing neurons in the mouse brain. J Comp Neurol, 519(10), 1847–1866. https://doi.org/10.1002/cne.22603

Moore, R. Y., Weis, R., & Moga, M. M. (2000). Efferent projections of the intergeniculate leaflet and the ventral lateral geniculate nucleus in the rat. J Comp Neurol, 420(3), 398–418. https://doi.org/10.1002/(sici)1096-9861(20000508)420:3<398::aid-cne9>3.0.co;2-9

Mu, D., Deng, J., Liu, K. F., Wu, Z. Y., Shi, Y. F., Guo, W. M., . . . Sun, Y. G. (2017). A central neural circuit for itch sensation. Science, 357(6352), 695–699. https://doi.org/10.1126/science.aaf4918

Nakamura, K., & Morrison, S. F. (2008). A thermosensory pathway that controls body temperature. Nat Neurosci, 11(1), 62–71. https://doi.org/10.1038/nn2027

Nakamura, K., & Morrison, S. F. (2010). A thermosensory pathway mediating heat-defense responses. Proc Natl Acad Sci U S A, 107(19),8848–8853. https://doi.org/10.1073/pnas.0913358107

Negishi, K., Payant, M. A., Schumacker, K. S., Wittmann, G., Butler, R. M., Lechan, R. M., . . . Chee, M. J. (2020). Distributions of hypothalamic neuron populations coexpressing tyrosine hydroxylase and the vesicular GABA transporter in the mouse. J Comp Neurol, 528(11), 1833–1855. https://doi.org/10.1002/cne.24857

Oishi, M., Kushikata, T., Niwa, H., Yakoshi, C., Ogasawara, C., Calo, G., . . . Hirota, K. (2014). Endogenous neuropeptide S tone influences sleep-wake rhythm in rats. Neurosci Lett, 581, 94–97. https://doi.org/10.1016/j.neulet.2014.08.031

Okamura, N., Hashimoto, K., Iyo, M., Shimizu, E., Dempfle, A., Friedel, S., & Reinscheid, R. K. (2007). Gender-specific association of a functional coding polymorphism in the Neuropeptide S receptor gene with panic disorder but not with schizophrenia or attention-deficit/hyperactivity disorder. Prog Neuropsychopharmacol Biol Psychiatry, 31(7), 1444–1448. https://doi.org/10.1016/j.pnpbp.2007.06.026

Opland, D., Sutton, A., Woodworth, H., Brown, J., Bugescu, R., Garcia, A., . . . Leinninger, G. (2013). Loss of neurotensin receptor-1 disrupts the control of the mesolimbic dopamine system by leptin and promotes hedonic feeding and obesity. Mol Metab, 2(4), 423–434. https://doi.org/10.1016/j.molmet.2013.07.008

Palmiter, R. D. (2018). The Parabrachial Nucleus: CGRP Neurons Function as a General Alarm. Trends Neurosci, 41(5), 280–293. https://doi.org/10.1016/j.tins.2018.03.007

Pape, H. C., Jungling, K., Seidenbecher, T., Lesting, J., & Reinscheid, R. K. (2010). Neuropeptide S: a transmitter system in the brain regulating fear and anxiety. Neuropharmacology, 58(1), 29–34. https://doi.org/10.1016/j.neuropharm.2009.06.001

Pauli, J. L., Chen, J. Y., Basiri, M. L., Park, S., Carter, M. E., Sanz, E., . . . Palmiter, R. D. (2022). Molecular and Anatomical Characterization of Parabrachial Neurons and Their Axonal Projections. bioRxiv.

Paxinos, G., & Franklin, K. B. J. (2013). Paxinos and Franklin’s The mouse brain in stereotaxic coordinates (Fourth edition. ed.). Academic Press, an imprint of Elsevier.

Paxinos, G., & Watson, C. (2007). The rat brain in stereotaxic coordinates (6th ed.). Academic Press/Elsevier.

Peng, Y. L., Han, R. W., Chang, M., Zhang, L., Zhang, R. S., Li, W., . . . Wang, R. (2010). Central Neuropeptide S inhibits food intake in mice through activation of Neuropeptide S receptor. Peptides, 31(12), 2259–2263. https://doi.org/10.1016/j.peptides.2010.08.015

Peng, Y. L., Zhang, J. N., Chang, M., Li, W., Han, R. W., & Wang, R. (2010). Effects of central neuropeptide S in the mouse formalin test. Peptides, 31(10), 1878–1883. https://doi.org/10.1016/j.peptides.2010.06.027

Petreanu, L., Huber, D., Sobczyk, A., & Svoboda, K. (2007). Channelrhodopsin-2-assisted circuit mapping of long-range callosal projections. Nat Neurosci, 10(5), 663–668. https://doi.org/10.1038/nn1891

Rizzi, A., Vergura, R., Marzola, G., Ruzza, C., Guerrini, R., Salvadori, S., . . . Calo, G. (2008). Neuropeptide S is a stimulatory anxiolytic agent: a behavioural study in mice. Br J Pharmacol, 154(2), 471–479. https://doi.org/10.1038/bjp.2008.96

Ruzza, C., Pulga, A., Rizzi, A., Marzola, G., Guerrini, R., & Calo’, G. (2012). Behavioural phenotypic characterization of CD-1 mice lacking the neuropeptide S receptor. Neuropharmacology, 62(5-6), 1999–2009. https://doi.org/10.1016/j.neuropharm.2011.12.036

Saldana, E., Vinuela, A., Marshall, A. F., Fitzpatrick, D. C., & Aparicio, M. A. (2007). The TLC: a novel auditory nucleus of the mammalian brain. J Neurosci, 27(48), 13108–13116. https://doi.org/10.1523/JNEUROSCI.1892-07.2007

Saldaña, E., Aparicio, M. A., Fuentes-Santamaría, V., & Berrebi, A. S. (2009). Connections of the superior paraolivary nucleus of the rat: projections to the inferior colliculus. Neuroscience, 163(1), 372–387. https://doi.org/10.1016/j.neuroscience.2009.06.030

Saper, C. B., & Loewy, A. D. (1980). Efferent connections of the parabrachial nucleus in the rat. Brain Res, 197(2), 291–317.

Shin, J. W., Geerling, J. C., Stein, M. K., Miller, R. L., & Loewy, A. D. (2011). FoxP2 brainstem neurons project to sodium appetite regulatory sites. J Chem Neuroanat, 42(1), 1–23. https://doi.org/10.1016/j.jchemneu.2011.05.003

Smith, K. L., Patterson, M., Dhillo, W. S., Patel, S. R., Semjonous, N. M., Gardiner, J. V., . . . Bloom, S. R. (2006). Neuropeptide S stimulates the hypothalamo-pituitary-adrenal axis and inhibits food intake. Endocrinology, 147(7), 3510–3518. https://doi.org/10.1210/en.2005-1280

Sun, J., Yuan, Y., Wu, X., Liu, A., Wang, J., Yang, S., . . . Huang, J. (2022). Excitatory SST neurons in the medial paralemniscal nucleus control repetitive self-grooming and encode reward. Neuron, 110(20), 3356–3373 e3358. https://doi.org/10.1016/j.neuron.2022.08.010

Varga, T., Palkovits, M., Usdin, T. B., & Dobolyi, A. (2008). The medial paralemniscal nucleus and its afferent neuronal connections in rat. J Comp Neurol, 511(2), 221–237. https://doi.org/10.1002/cne.21829

Viñuela, A., Aparicio, M. A., Berrebi, A. S., & Saldaña, E. (2011). Connections of the Superior Paraolivary Nucleus of the Rat: II. Reciprocal Connections with the Tectal Longitudinal Column. Front Neuroanat, 5, 1. https://doi.org/10.3389/fnana.2011.00001

Vrang, N., Mrosovsky, N., & Mikkelsen, J. D. (2003). Afferent projections to the hamster intergeniculate leaflet demonstrated by retrograde and anterograde tracing. Brain Res Bull, 59(4), 267–288. https://doi.org/10.1016/s0361-9230(02)00875-4

Vujovic, N., Gooley, J. J., Jhou, T. C., & Saper, C. B. (2015). Projections from the subparaventricular zone define four channels of output from the circadian timing system. J Comp Neurol, 523(18), 2714–2737. https://doi.org/10.1002/cne.23812

Walker, D. L., Toufexis, D. J., & Davis, M. (2003). Role of the bed nucleus of the stria terminalis versus the amygdala in fear, stress, and anxiety. Eur J Pharmacol, 463(1-3), 199–216. https://doi.org/10.1016/s0014-2999(03)01282-2

Wang, J., Palkovits, M., Usdin, T. B., & Dobolyi, A. (2006). Afferent connections of the subparafascicular area in rat. Neuroscience, 138(1), 197–220. https://doi.org/10.1016/j.neuroscience.2005.11.010

Wang, Q., Ding, S. L., Li, Y., Royall, J., Feng, D., Lesnar, P., . . . Ng, L. (2020). The Allen Mouse Brain Common Coordinate Framework: A 3D Reference Atlas. Cell, 181(4), 936–953.e920. https://doi.org/10.1016/j.cell.2020.04.007

Wouterlood, F. G., & Jorritsma-Byham, B. (1993). The anterograde neuroanatomical tracer biotinylated dextran-amine: comparison with the tracer Phaseolus vulgaris-leucoagglutinin in preparations for electron microscopy. J Neurosci Methods, 48(1-2), 75–87. https://doi.org/10.1016/s0165-0270(05)80009-3

Xing, L., Shi, G., Mostovoy, Y., Gentry, N. W., Fan, Z., McMahon, T. B., . . . Fu, Y. H. (2019). Mutant neuropeptide S receptor reduces sleep duration with preserved memory consolidation. Sci Transl Med, 11(514). https://doi.org/10.1126/scitranslmed.aax2014

Xu, Y. L., Gall, C. M., Jackson, V. R., Civelli, O., & Reinscheid, R. K. (2007). Distribution of neuropeptide S receptor mRNA and neurochemical characteristics of neuropeptide S-expressing neurons in the rat brain. J Comp Neurol, 500(1), 84–102. https://doi.org/10.1002/cne.21159

Xu, Y. L., Reinscheid, R. K., Huitron-Resendiz, S., Clark, S. D., Wang, Z., Lin, S. H., . . . Civelli, O. (2004). Neuropeptide S: a neuropeptide promoting arousal and anxiolytic-like effects. Neuron, 43(4), 487–497. https://doi.org/10.1016/j.neuron.2004.08.005

Yassa, M. A., Hazlett, R. L., Stark, C. E., & Hoehn-Saric, R. (2012). Functional MRI of the amygdala and bed nucleus of the stria terminalis during conditions of uncertainty in generalized anxiety disorder. J Psychiatr Res, 46(8), 1045–1052. https://doi.org/10.1016/j.jpsychires.2012.04.013

Zhu, H., Mingler, M. K., McBride, M. L., Murphy, A. J., Valenzuela, D. M., Yancopoulos, G. D., . . . Rothenberg, M. E. (2010). Abnormal response to stress and impaired NPS-induced hyperlocomotion, anxiolytic effect and corticosterone increase in mice lacking NPSR1. Psychoneuroendocrinology, 35(8), 1119–1132. https://doi.org/10.1016/j.psyneuen.2010.01.012

Zhu, Y. B., Wang, Y., Hua, X. X., Xu, L., Liu, M. Z., Zhang, R., . . . Mu, D. (2022). PBN-PVT projections modulate negative affective states in mice. Elife, 11. https://doi.org/10.7554/eLife.68372

