## Supplementary figures and images for "Efferent projections of *Nps*-expressing neurons in the parabrachial region"

### Supplemental Figure 1 (full rostrocaudal injection sites, Syp-mCherry NiDAB labeling with Nissl counterstain)

5429

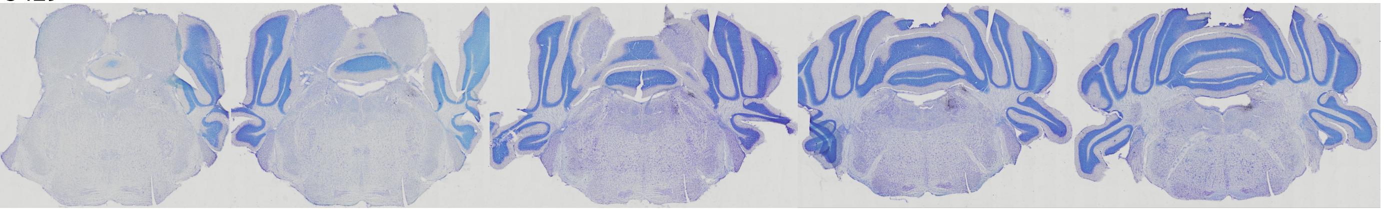

5432

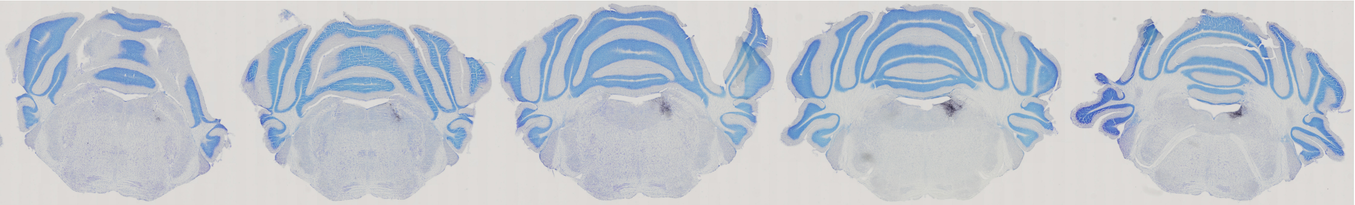

5495

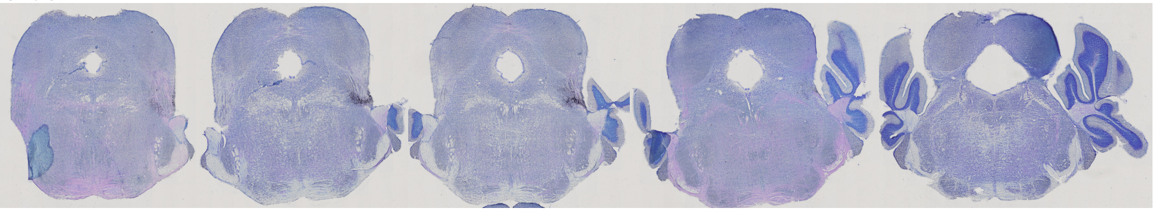

5568

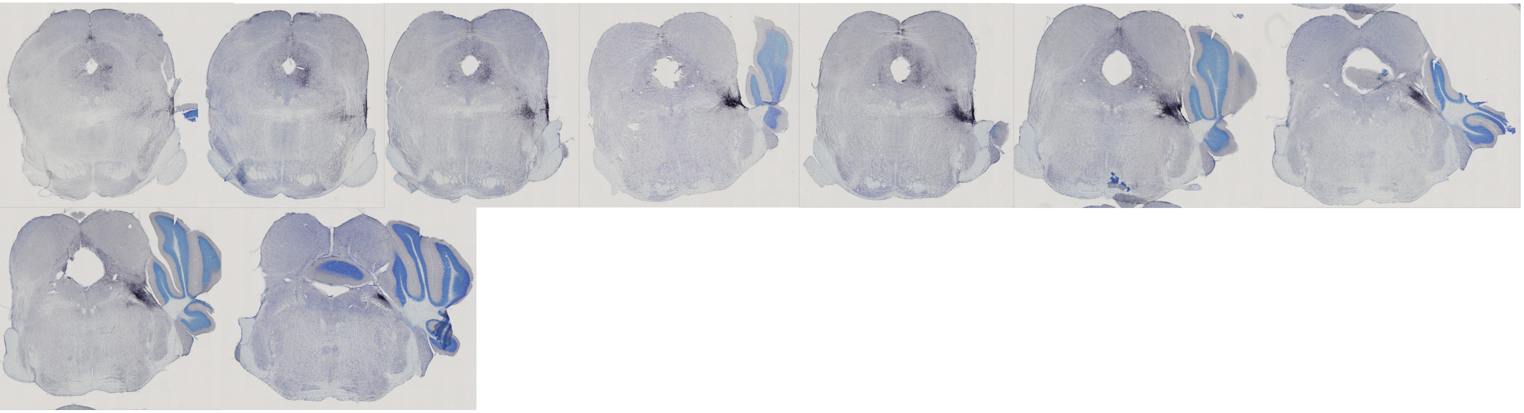

5569

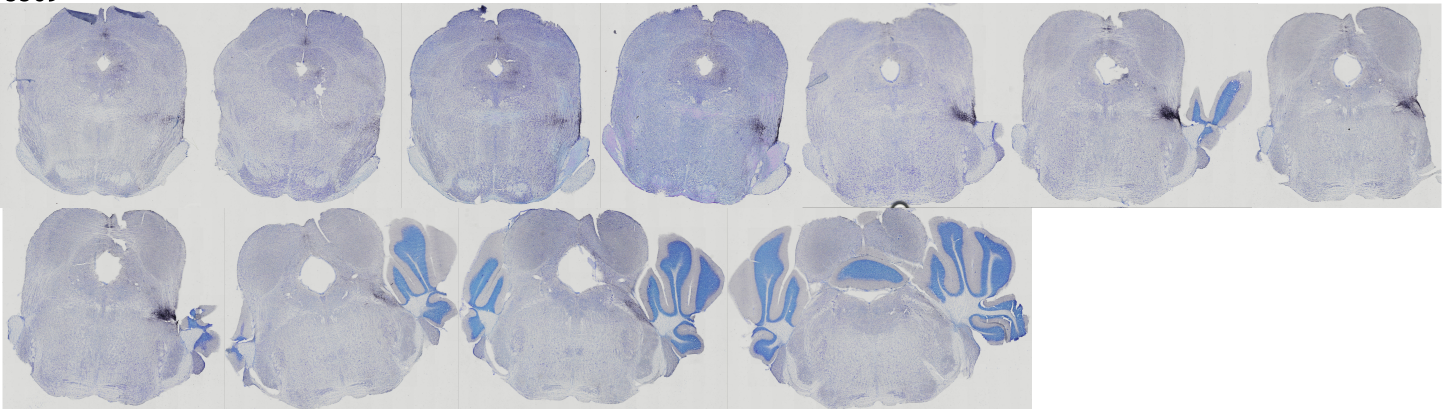
